# Novel inhibitors of *E. coli* lipoprotein diacylglyceryl transferase are insensitive to resistance caused by *lpp* deletion

**DOI:** 10.1101/2020.10.04.325589

**Authors:** Jingyu Diao, Rie Komura, Tatsuya Sano, Homer Pantua, Kelly M. Storek, Hiroko Inaba, Haruhiko Ogawa, Cameron L. Noland, Yutian Peng, Susan L. Gloor, Donghong Yan, Jing Kang, Anand Kumar Katakam, Nicholas N. Nickerson, Cary D. Austin, Jeremy Murray, Steven T. Rutherford, Mike Reichelt, Yiming Xu, Min Xu, Hayato Yanagida, Junichi Nishikawa, Patrick C Reid, Christian N. Cunningham, Sharookh B. Kapadia

## Abstract

Lipoprotein diacylglyceryl transferase (Lgt) catalyzes the first step in the biogenesis of Gram-negative bacterial lipoproteins which play crucial roles in bacterial growth and pathogenesis. We demonstrate that Lgt depletion in a clinical uropathogenic *Escherichia coli* strain leads to permeabilization of the outer membrane and increased sensitivity to serum killing and antibiotics. Importantly, we identify the first ever described Lgt inhibitors that potently inhibit Lgt biochemical activity *in vitro* and are bactericidal against wild-type *Acinetobacter baumannii* and *E. coli* strains. Unlike inhibition of other steps in lipoprotein biosynthesis, deletion of the major outer membrane lipoprotein, *lpp*, is not sufficient to rescue growth after Lgt depletion or provide resistance to Lgt inhibitors. Our data validate Lgt as a novel druggable antibacterial target and suggest that inhibition of Lgt may not be sensitive to one of the most common resistance mechanisms that invalidate inhibitors of downstream steps of bacterial lipoprotein biosynthesis and transport.

## Introduction

The cell envelope of a typical Gram-negative bacterium consists of two membranes: a phospholipid inner membrane (IM) and an asymmetrical outer membrane (OM), the latter of which is composed of a phospholipid inner leaflet and a lipopolysaccharide (LPS) outer leaflet. The IM and OM are separated by the periplasm, which contains a peptidoglycan (PG) cell wall (reviewed in detail in (Silhavy, Kahne, & Walker, 2010)). *E. coli* encodes >90 lipoproteins, many of which are localized to the inner leaflet of the OM, but can also be exposed on the bacterial cell surface (Cowles, Li, Semmelhack, Cristea, & Silhavy, 2011; Wilson & Bernstein, 2015). Bacterial lipoproteins play critical roles in adhesion, nutrient uptake, antibiotic resistance, virulence, invasion and immune evasion (Kovacs-Simon, Titball, & Michell, 2011), making the lipoprotein biosynthetic and transport pathways attractive targets for novel antibacterial drug discovery.

Lipoprotein biosynthesis in Gram-negative bacteria is mediated by three IM localized enzymes: Lgt, LspA and Lnt (Figure 1). All preprolipoproteins contain a signal peptide followed by a conserved four amino acid sequence, [LVI][ASTVI][GAS]C, also known as a lipobox (Schlesinger, 1992), and are secreted through the IM via the Sec or Tat pathways. After secretion through the IM, Lgt catalyzes the attachment of a diacylglyceryl moiety from phosphatidylglycerol to the thiol group of the conserved +1 position cysteine via a thioether bond (Sankaran & Wu, 1994). The second enzyme, prolipoprotein signal peptidase (LspA), is an aspartyl endopeptidase which cleaves off the signal peptide N-terminal of the conserved diacylated +1 cysteine (M. Tokunaga, Tokunaga, & Wu, 1982), and is the molecular target of the Gram-negative-specific natural-product antibiotics globomycin and myxovirescin (Dev, Harvey, & Ray, 1985; Gerth, Irschik, Reichenbach, & Trowitzsch, 1982; Olatunji et al., 2020; Xiao, Gerth, Müller, & Wall, 2012). In Gram-negative and high-GC Gram-positive bacteria, a third enzyme, lipoprotein N-acyl transferase (Lnt), catalyzes the addition of a third acyl chain to the amino group of the N-terminal cysteine via an amide linkage. Mature triacylated lipoproteins destined for the OM are extracted from the IM by the LolCDE ATP-binding cassette (ABC) transporter and transported to the OM via a periplasmic chaperone protein LolA and an OM lipoprotein LolB (Narita, 2011; Narita & Tokuda, 2010) (Figure 1).

**Figure 1:**
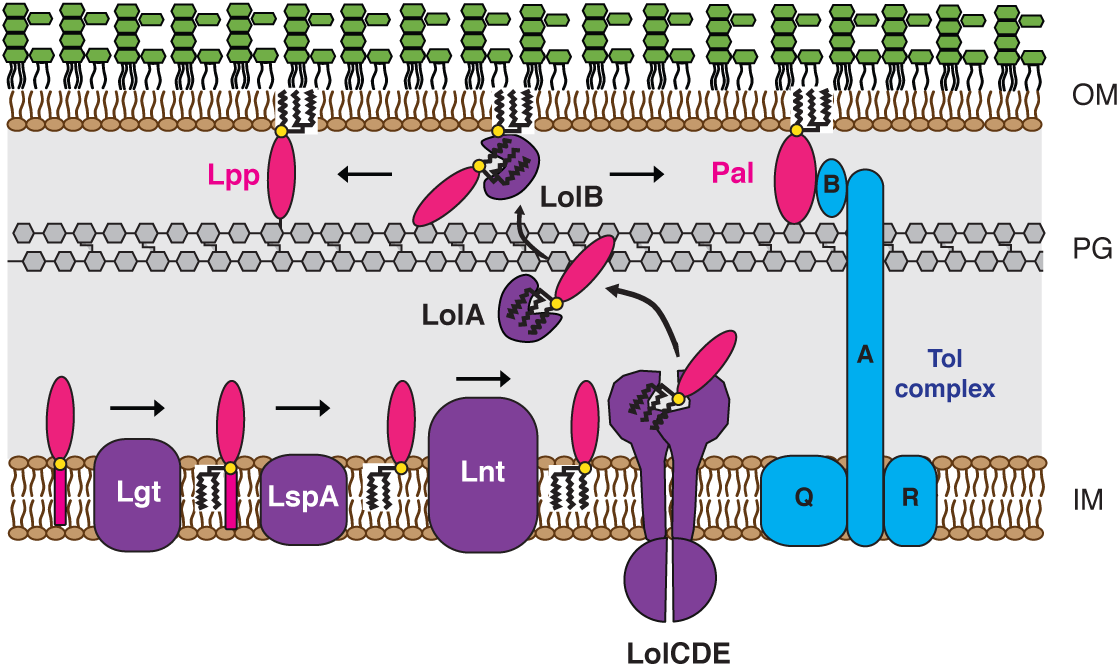
Lipoprotein biosynthesis and transport in Gram-negative bacteria. Prolipoprotein substrates translocate through the IM via the Sec or Tat pathway and are sequentially modified by Lgt, LspA and Lnt. Triacylated lipoproteins that are destined for the OM are recognized by the Lol system (LolABCDE) and transported to the OM. Lpp and Pal are two OM lipoproteins that tether the OM to the PG layer. Pal also binds to TolB, which also can interact with Lpp and OmpA, an OM *β*-barrel protein that can also associate with PG (not shown).

Two OM lipoproteins, Lpp (also known as Murein lipoprotein or Braun’s lipoprotein) and Pal (peptidoglycan-associated lipoprotein), mediate tethering of the PG layer to the OM in *E. coli*. Lpp is a small ∼8 kDa lipoprotein that is the most abundant OM protein in *E. coli* (∼500,000 molecules per cell) and a third of all Lpp is covalently linked to PG (Cowles et al., 2011; Neidhardt, 1996). *E. coli* mutants deficient in Lpp exhibit increased OM permeability, leakage of periplasmic components, increased outer membrane vesicle (OMV) release and increased sensitivity to complement-mediated lysis (Diao et al., 2017; H. Suzuki et al., 1978; Yem & Wu, 1978). Mislocalization and accumulation of PG-linked Lpp in the inner membrane upon inhibition of LspA (Xiao et al., 2012; Zwiebel, Inukai, Nakamura, & Inouye, 1981) and LolCDE (McLeod et al., 2015; Nickerson et al., 2018) is believed to lead to bacterial cell death (Narita & Tokuda, 2011; Robichon, Vidal-Ingigliardi, & Pugsley, 2005; Yakushi, Tajima, Matsuyama, & Tokuda, 1997a). In addition to Lpp, Pal binds PG and interacts with OmpA, Lpp and the Tol complex, and is crucial for maintaining OM integrity in *E. coli* (Cascales, Bernadac, Gavioli, Lazzaroni, & Lloubes, 2002; Clavel, Germon, Vianney, Portalier, & Lazzaroni, 1998; Leduc, Ishidate, Shakibai, & Rothfield, 1992; Mizuno, 1979). While non-natural product inhibitors of LspA and LolCDE have been previously discovered (Kitamura, Owensby, Wall, & Wolan, 2018; McLeod et al., 2015), no inhibitors of the first committed step in bacterial lipoprotein biosynthesis have been described. Since many natural product antibiotics, including those that inhibit LspA, are cyclic (Igarashi, 2019; Rossiter, Fletcher, & Wuest, 2017), we screened a macrocyclic peptide library to identify Lgt inhibitors. In this study, we identify and characterize the first inhibitors of Lgt that inhibit growth of wild-type *E. coli* and *A. baumannii* strains in addition to other OM-permeabilized Gram-negative species. We demonstrate that, unlike inhibitors of LspA and LolCDE, treatment with Lgt inhibitors does not lead to the significant accumulation of PG-linked Lpp forms in the IM and as such, are not sensitive to resistance mediated by deletion of *lpp*.

## Results

### Modest depletion of Lgt leads to increased OM permeability and loss of bacterial viability is not rescued by deletion of *lpp*

Previous investigations into the role of Lgt in *E. coli* have focused on laboratory strains, specifically those lacking the O-antigen of LPS. Here, we engineered the uropathogenic *E. coli* clinical isolate CFT073 so that the only copy of *lgt* was under control of an arabinose-inducible promoter (CFT073*Δlgt*), and hence requires arabinose for Lgt expression. As expected, genetic depletion of Lgt was lethal *in vitro* and growth was rescued after complementation with *E. coli lgt* (Figure 2a). *thyA*, the gene that encodes thymidylate synthase, is downstream of *lgt* and its ribosome binding site overlaps with the *lgt* stop codon. We confirmed that *thyA* expression, which is regulated by transcription from the *lgt* promoter and translational coupling (Gan et al., 1995), was unchanged after Lgt depletion (Figure 2-figure supplement 1a).

**Figure 2:**
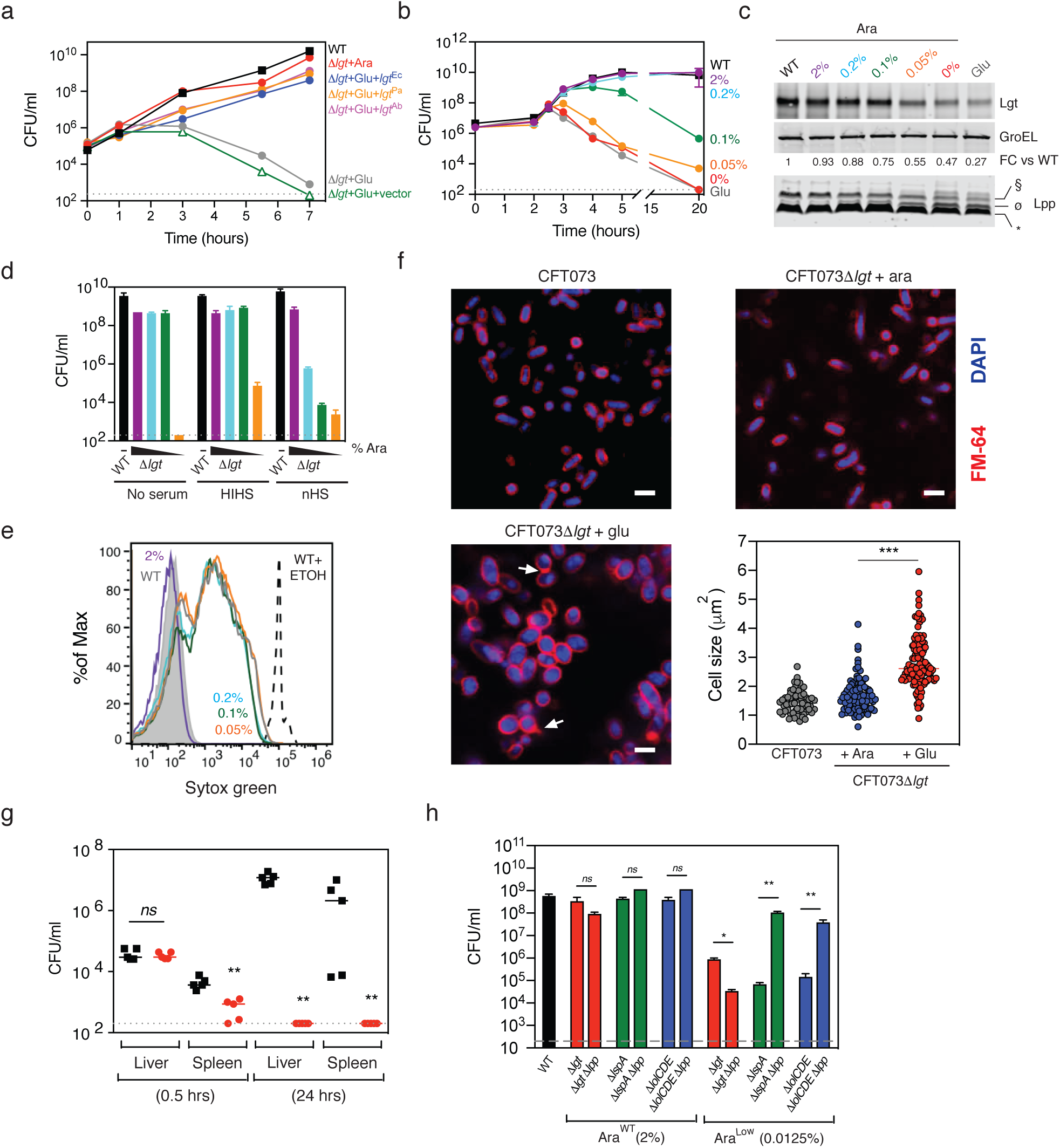
Lgt is essential for *in vitro* growth, membrane integrity, serum resistance and virulence. **(a)** CFT073*Δlgt* cells were grown in the presence of 4% arabinose (red circles) or 0.2% glucose (grey circles) and CFUs were enumerated over 7 hours post treatment. CFT073*Δlgt* cultured in the presence of 0.2% glucose were complemented with empty pLMG18 plasmid (open green triangles) or pLMG18 plasmids expressing *lgt* from *E. coli* (blue circles), *A. baumannii* (magenta circles) or *P. aeruginosa* (orange circles). The grey dashed line represents the limit of detection (200 CFU/ml) of the experiment. Data are representative of two independent experiments each performed in duplicate. **(b-c)** A modest ∼25% reduction in Lgt levels results in a significant loss in viability over time with a concurrent accumulation of the unmodified pro-Lpp (ø, UPLP). CFT073*Δlgt* cells were treated with a range of arabinose concentrations and CFUs were enumerated over 20 hours. CFU growth data are representative of two independent experiments each performed in duplicate. Western blot analysis for expression of Lgt and Lpp was performed using WT CFT073 and CFT073*Δlgt* total cell lysates harvested at 3 hours post arabinose treatment. To quantitate Lgt expression levels, Lgt levels were normalized to GroEL and quantitated as fold change relative to WT CFT073 (FC vs WT). Lpp forms are denoted as follows: * = triacylated free Lpp; § = PG-linked diacylglyceryl pro-Lpp (DGPLP); ø = unmodified pro-Lpp (UPLP). Data are representative of two independent experiments. **(d)** Lgt depletion leads to increased serum sensitivity. WT CFT073 and CFT073*Δlgt* cells grown in the presence of a range of arabinose concentrations (2% = magenta; 0.2% = light blue; 0.1% = green and 0.05% = orange) were incubated with 50% normal human serum (nHS), heat inactivated human serum (HIHS) or medium (no serum) for 1 hour and CFUs were enumerated. Data are representative of at least three independent experiments each performed in duplicate. **(e)** Lgt depletion leads to increased OM permeability. WT CFT073 and CFT073*Δlgt* cells were incubated with the same range of arabinose concentrations as in Figure 2d and incubated with the nucleic acid dye, SYTOX Green, and flow cytometry was performed to determine level of dye incorporation. While SYTOX Green does not efficiently incorporate in bacterial cells with an intact OM (CFT073*Δlgt* treated with 2% arabinose, magenta), SYTOX Green incorporation in bacterial cells increases after Lgt depletion. Intact CFT073 (WT, grey) or CFT073 treated with 70% ethanol (WT+ETOH, black), which permeabilizes the cells, were used as controls. Data are representative of two independent experiments. **(f)** Lgt depletion results in a globular cellular phenotype and membrane blebbing. WT CFT073 or CFT073*Δlgt* cells were grown in either arabinose or glucose for 4 hours, fixed and incubated with FM-64 dye (red) and DAPI (blue) to detect OM and nucleic acids, respectively. Cells were visualized by confocal microscopy. Arrows represent membrane blebs. Scale bars represent 1 µm. Quantitation of cell size was performed using ImageJ software. **(g)** Lgt depletion leads to significant attenuation in virulence. Intravenous infection of neutropenic A/J mice with WT CFT073 (black) or CFT073*Δlgt* (red) cells. At 0.5 hours and 24 hours post-infection, bacterial burden in the liver and spleen were enumerated. Overall *p*-value for the ANOVA is *p* <0.0001. Pairwise comparisons were analyzed using unpaired Mann Whitney test (** *p* = 0.0079). The grey dashed line represents the limit of detection (200 CFU/ml) for this experiment. **(h)** Deletion of *lpp* does not rescue growth after Lgt depletion. *E. coli* MG1655 (WT, black), or inducible deletion strains for *lgt* (*Δlgt*, red), *lspA* (*ΔlspA, green)* and *lolCDE* (*ΔlolCDE*, blue) that either contained *lpp* or had *lpp* deleted were grown in conditions that allowed for normal growth (Ara^WT^, 2% arabinose) or decreased growth (Ara^Low^, 0.0125% arabinose) and CFUs at were enumerated at 5 hours post treatment. Data are representative of two independent experiments each performed in duplicate (*ns* = not significant, *p < 0.05, **p < 0.01).

Complementation with *lgt* from *Pseudomonas aeruginosa* PA14 or *A. baumannii* ATCC 17978 (51.6% and 48.6% sequence identity, respectively) was able to rescue viability (Figure 2a and Figure 2-figure supplement 1b). Overexpression of the *E. coli* genes encoding the downstream enzymes in lipoprotein biosynthesis (LspA, Lnt) and transport (LolCDE) did not rescue growth of CFT073*Δlgt* in spite of detectable levels of LspA, Lnt and LolCDE (Figure 2-figure supplement 1c - g). While depletion of ∼25% of Lgt was sufficient for bactericidal activity (Figure 2b and 2c), CFT073*Δlgt* cells expressing as high as ∼90% of normal levels of Lgt were significantly more sensitive to complement-mediated killing of the normally serum-resistant *E. coli* CFT073 and showed increased incorporation of SYTOX Green, a dye that normally does not penetrate an intact OM (Figure 2c-e). Depletion of Lgt also resulted in an expected increase in cell size (Figure 2f) and an Lpp-dependent IM contraction due to osmotic stress (Figure 2-figure supplement 2), as previously reported (Inukai et al., 1978a; Inukai, Nakajima, Osawa, Haneishi, & Arai, 1978b; Rojas et al., 2018). Consistent with these results, partial depletion of Lgt that still allowed for normal growth *in vitro* led to increased sensitivity to antibiotics that are normally excluded by the impermeable Gram-negative OM (Table 1). Depletion of Lgt also resulted in significant attenuation in a mouse *E. coli* bacteremic infection model (Figure 2g). Cumulatively, these data suggest that Lgt could be a good antibiotic target since partial inhibition of Lgt may be sufficient to lead to significant attenuation in growth and cellular morphology.

**Table 1:**
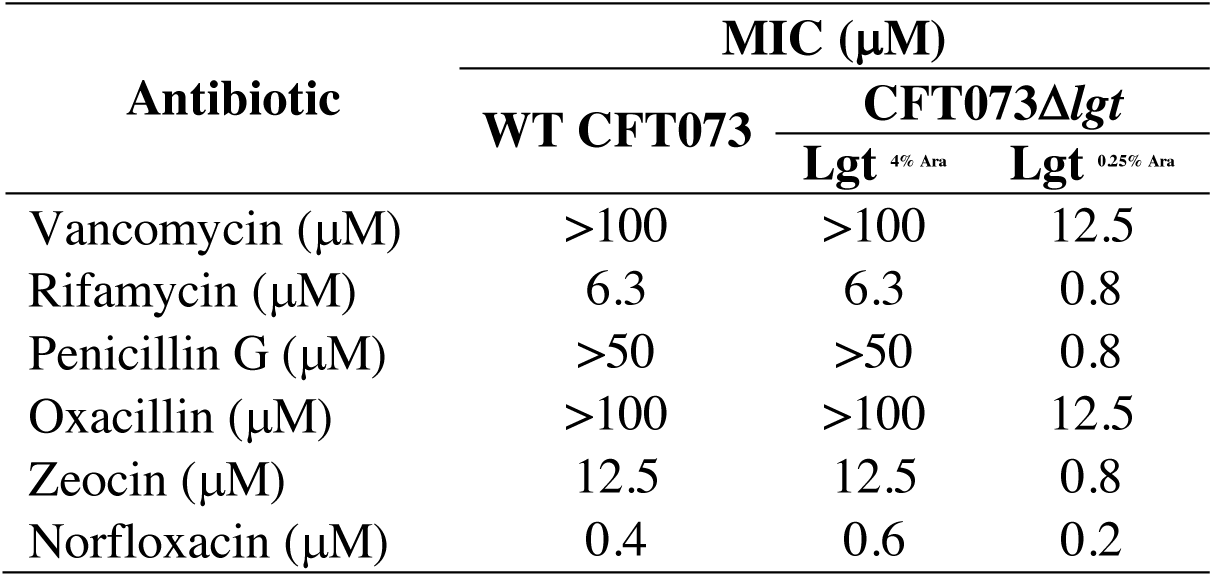
Antibiotic sensitivity of WT CFT073 versus CFT073*Δlgt* cells expressing wild-type (4% Ara) or low (0.25% Ara) levels of Lgt

Bactericidal activity of LspA and LolCDE inhibitors are sensitive to deletion of the gene encoding the major OM lipoprotein, Lpp (McLeod et al., 2015; Zwiebel et al., 1981). To determine if Lpp played a role in bacterial cell death after Lgt depletion, we constructed a *lgt* inducible deletion strain in *E. coli* MG1655 with and without *lpp* (MG1655Δ*lgt* and MG1655Δ*lgt*Δ*lpp*) and compared growth of these strains to *lspA* and *lolCDE* inducible deletion strains in the same backgrounds. Expectedly, *lpp* deletion rescued the growth of the *lspA* and *lolCDE* inducible deletions strains after depletion of LspA and LolCDE, respectively (Figure 2h). In contrast to LspA and LolCDE depletion, the *lpp* mutant was more sensitive to Lgt depletion leading to a greater loss of colony forming units (CFU) compared to that detected after Lgt depletion in cells expressing *lpp*.

Since the loss of *lpp* is a primary mechanism of resistance to inhibitors of LspA and LolCDE thereby complicating their potential as antibacterial targets, identification of Lgt inhibitors would uncover further biological understanding of this essential pathway, and potentially serve as better starting chemical matter to develop novel antibiotics targeting lipoprotein biosynthesis that are not sensitive to resistance mediated by *lpp* deletion.

### Identification and characterization of macrocyclic peptide inhibitors of Lgt

Many natural products or their derivatives account for a significant number of launched drugs and sine many of them are cyclic in nature (Igarashi, 2019), we initially screened a macrocyclic peptide library to identify specific and high affinity binders of Lgt. A genetically reprogrammed *in vitro* translation system combined with mRNA affinity selection methods was used to generate large macrocycle peptide libraries with sizes varying from 8-14 amino acids in length (Goto, Katoh, & Suga, 2011; Ishizawa, Kawakami, Reid, & Murakami, 2013; Kashiwagi, Reid, & Inc, 2013) (Figure 3a). The variable sequence (6-12 amino acids) of the macrocycle libraries encoded the random incorporation of 11 natural amino acids (Ser, Tyr, Trp, Leu, Pro, His, Arg, Asn, Val, Asp, and Gly) and 5 non-natural amino acids (Figure 3b). The screening of the libraries is schematically depicted in Figure 3c. Lgt-biotin was solubilized in 0.02% n-Dodecyl β-D-maltoside (DDM), immobilized on streptavidin magnetic beads and incubated with the macrocyclic library. Iterative rounds of affinity selection were performed to identify Lgt-binding macrocycles. After five rounds of enrichment, two additional rounds of off-rate selections were performed by increasing the wash stringency before high affinity binders were eluted. Hit macrocycles were identified using next generation sequencing on the last four rounds of selection followed by a frequency analysis calculation. Three macrocycles were identified from these screens, **508**, **692**, and **693** (Figure 3-figure supplement 1), with a frequency enrichment in the final round of selection of 4.1%, 19.4%, and 10.1%, respectively, as measured by NGS. **508** contains 8 amino acids with a molecular weight (MW) of 1264.49 Da. **692** and **693** each contain 7 amino acids and are related to one another with a charge swap at position 2, and have MWs of 1428.66 and 1259.55 respectively. The calculated LogPs (cLogP), which is the logarithm of the compounds partition coefficient between n-octanol and water and a measure of a molecule’s hydrophilicity, were 4, 1.8 and 1.7 for **508**, **692** and **693**, respectively. **692** was synthesized with a Gly off of the C-terminus and renamed G9066 (Figure 3d). During re-synthesis, both **508** and **693** were synthesized with a Gly-Lys-Lys tail off of the C-terminus to aide in solubility of these macrocycles and were renamed G2823 and G2824, respectively (Figure 3d).

**Figure 3:**
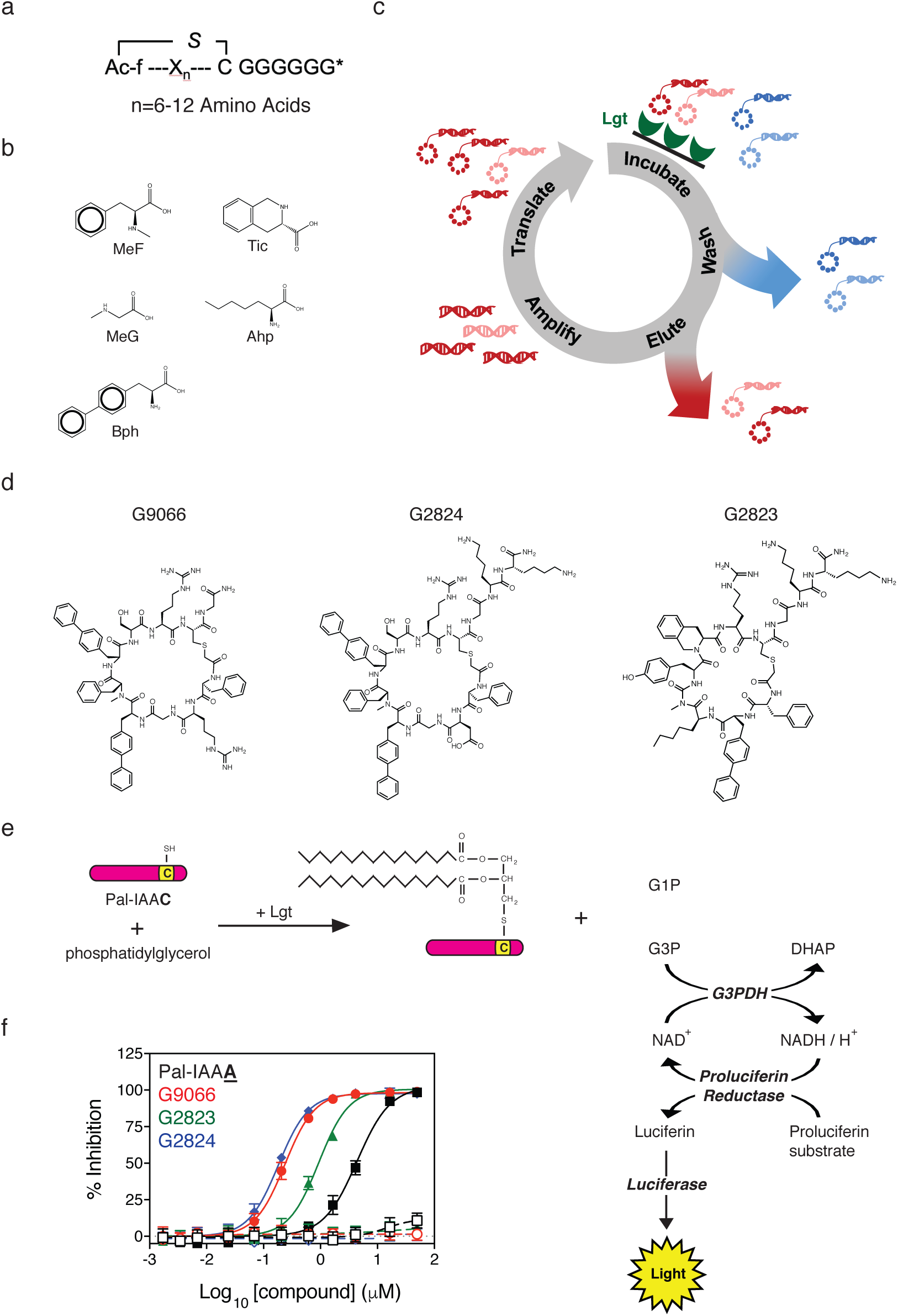
Identification of Lgt inhibitors. **(a)** Representation of macrocycle peptide libraries varying in size from 8-14 amino acids in length. The variable region (X_n_) of the macrocycle libraries was encoded to allow the random incorporation of 11 natural amino acids and 5 non-natural amino acids. **(b)** The 5 non-natural amino acids used in the generation of the libraries were N-*α*-Methyl-L-phenylalanine (MeF), N-*α*-Methyl-L-glycine (MeG, Sarcosine), (S)-2-Aminoheptanoic acid (Ahp), 4-Phenyl-L-phenylalanine (Bph) and (S)-1,2,3,4-Tetrahydroisoquinoline-3-carboxylic acid (Tic). **(c)** Schematic representation of affinity-based selections using recombinant Lgt-biotin immobilized on streptavidin magnetic beads. As discussed in the Methods, Lgt-DDM was incubated with the macrocycle library and Lgt binders were eluted, amplified and translated to generate new libraries enriched for Lgt binders. Iterative rounds of affinity selection and washing were performed against recombinant Lgt and macrocycles that bound to Lgt were identified using next generation sequencing. **(d)** Structure of the macrocyclic peptides G9066, G2823 and G2824 identified in this study. **(e)** Development of the *in vitro* Lgt biochemical assay. Lgt-DDM was incubated with phosphatidylglycerol and the Pal-IAAC peptide substrate derived from the Pal lipoprotein (MQLNKVLKGLMIALPVMAIAA<u>C</u>SSNKN) for 60 minutes at RT, as described in the Methods. After Lgt catalyzes the transfer of diacylglyceryl from phosphatidylglycerol to the Pal substrate (Pal-IAAC), glycerol-1-phosphophate (G1P) is released from phosphatidylglycerol. Given the phosphatidylglycerol substrate used in our biochemical assay contains a racemic glycerol moiety at the end of phosphatidyl group, both G1P and G3P are released. G3P is quantitatively converted to Dihydroxyacetone phosphate (DHAP) with concomitant formation of an equivalent amount of NADH by the action of glycerol 3-phosphate dehydrogenase (G3PDH). Newly formed NADH will in turn quantitatively react with proluciferin to generate equivalent amounts of luciferin, which ultimately results in luminescence by luciferase that is proportional to the amount of luciferin available. **(f)** Dose-dependent inhibition of Lgt biochemical activity. Lgt was incubated with phosphatidylglycerol and the Pal-IAAC substrate in the presence of absence of by G9066 (red), G2823 (green) or G2824 (blue). Luminescence values were normalized to DMSO controls (0% inhibition) and no enzyme controls (100% inhibition). As a control, we incubated the Lgt reactions with a mutant substrate peptide also derived from the Pal lipoprotein which has the conserved cysteine mutated to alanine (Pal-IAA**<u>A</u>**, black). While the Pal-IAA**<u>A</u>** peptide binds to Lgt, it cannot be modified by Lgt and acts as a non-modifiable, competitive peptide. Negative control reactions for each inhibitor were run in the absence of Lgt enzymes (open symbols). Data are representative of at least two independent experiments each performed in triplicate.

We then tested the ability of G9066, G2823 and G2824 to inhibit *E. coli* Lgt enzymatic activity *in vitro* by measuring the release of glycerol phosphate which is a by-product of the Lgt-catalyzed transfer of diacylglyceryl from phosphatidylglycerol to a peptide substrate via formation of a thioether bond. The peptide substrate was derived from the Pal lipoprotein (Pal-IAAC, where C is the conserved cysteine that is modified by Lgt). While glycerol-1-phosphate (G1P) is the expected by-product of the Lgt enzymatic activity (Sankaran & Wu, 1994), the phosphatidylglycerol substrate used in our biochemical assay contains a racemic glycerol moiety at the end of phosphatidyl group, and hence both G1P and glycerol-3-phosphate (G3P) are released from phosphatidylglycerol as Lgt catalyzes the reaction (Figure 3e). The detection of G3P is based on a coupled luciferase reaction which is described in more detail in the Methods and in Figure 3e. G9066, G2823 and G2824 potently inhibited Lgt biochemical activity (IC_50_=0.24 μM, 0.93 μM and 0.18 μM, respectively) (Figure 3f). In comparison, a mutant Pal peptide substrate with the conserved cysteine mutated to alanine (Pal-IAA**<u>A</u>**), which cannot get modified and acts as a Lgt-binding nonreactive, substrate-based competitive inhibitor, inhibited Lgt with an IC_50_=4.4 μM (Figure 3f). When tested against bacterial cells in minimal inhibitory concentration (MIC) growth assays, G9066 and G2824 inhibited growth of WT *A. baumannii* 19606 with a MIC = 37.5 μM. G2823 and G2824 inhibited *E. coli* MG1655 growth with a MIC = 50 μM (Table 2). OM permeabilization either genetically (*imp*4213 mutation) or chemically (EDTA treatment) of all Gram-negative strains, including *P. aeruginosa* PA14 and *A. baumannii* 19606, led to growth inhibition (Table 2). Interestingly, *lpp* deletion in either CFT073*imp*4213 or CFT073 treated with EDTA did not lead to increases in G9066, G2823 or G2824 MIC, unlike that seen with inhibitors of LspA and LolCDE (Table 2). In fact, *lpp* deletion in WT CFT073 cells led to a modest increase in G9066, G2823 and G2824 potency. G2823 and G2824 showed minimal non-specific activity against eukaryotic cells and the Gram-positive *Staphylococcus aureus* strain USA300, consistent with data demonstrating *lgt* is dispensable for Gram-positive bacterial growth *in vitro* (Stoll, Dengjel, Nerz, & Götz, 2005). In contrast, G9066 inhibited growth of USA300 to a greater extent suggesting G9066 may have additional targets or non-specific cellular effects. Given G9066 and G2824 are very similar, we decided to focus the remainder of this study on G2823 and G2824 (hereafter referred to as Lgti).

**Table 2:**
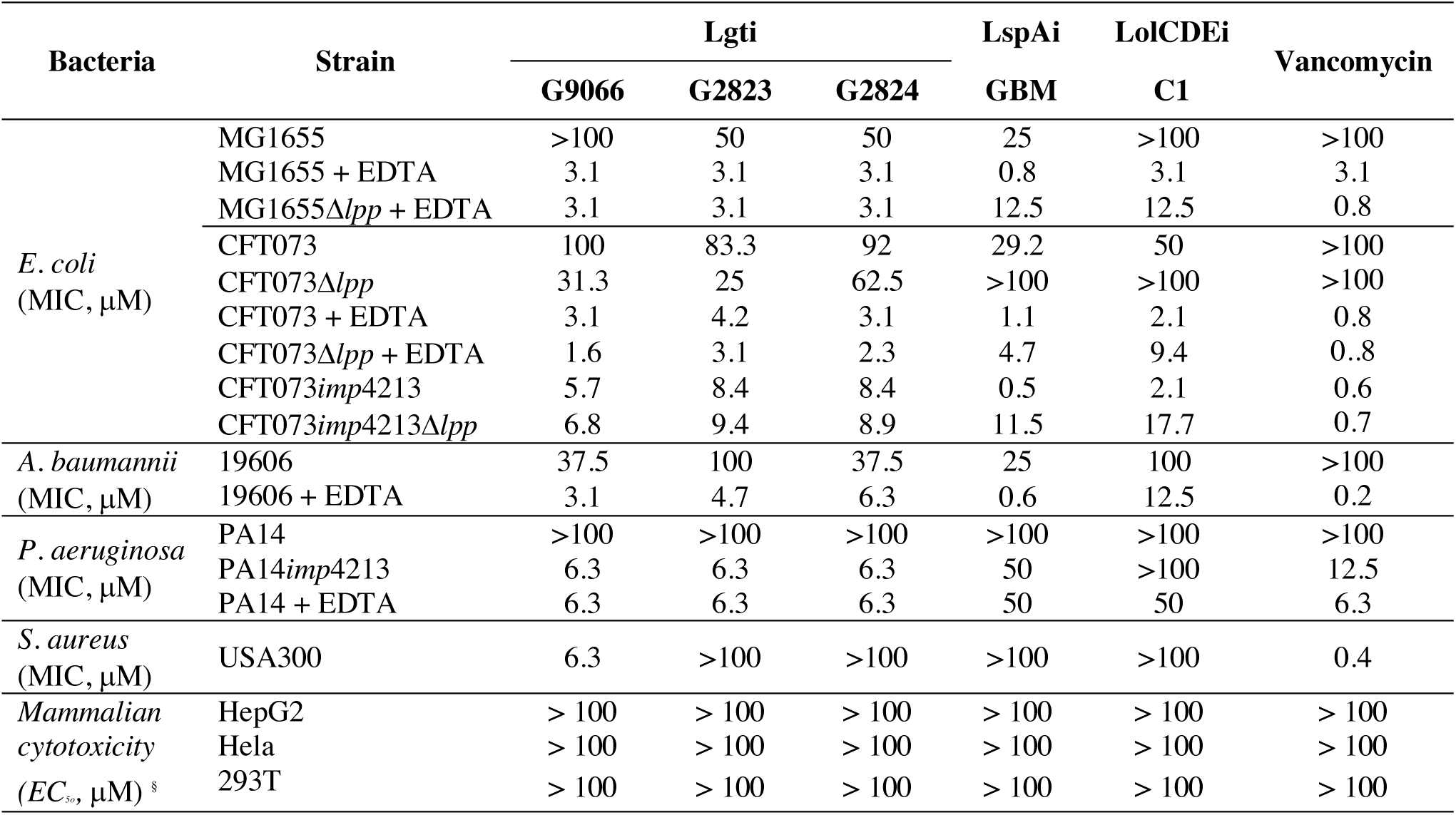
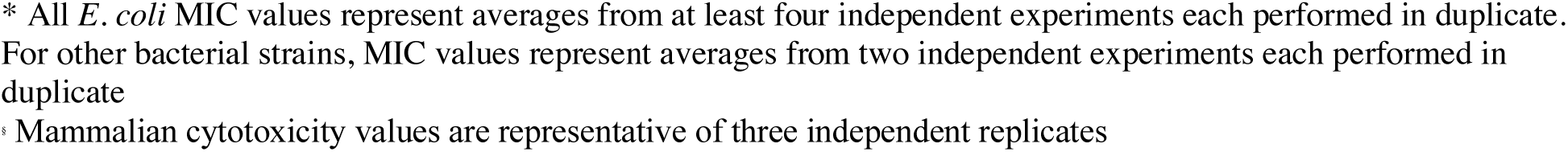
Growth inhibition of a panel of bacterial strains and eukaryotic cells by Lgti, LspAi and LolCDEi

### G2823 and G2824 specifically inhibit Lgt in *E. coli*

While the Lgti inhibited both Lgt enzymatic function and bacterial growth, it was unclear whether inhibition of bacterial cell growth was mediated by specific inhibition of Lgt function. We were unable to raise on-target resistant mutants to Lgti, and hence multiple experimental approaches were undertaken to determine if inhibition of bacterial growth was indeed Lgt-dependent. As the accumulation of Lpp intermediates detected by Western blot analyses has been successfully used to verify inhibition or deletion of specific enzymes involved in lipoprotein biosynthesis or transport (Narita & Tokuda, 2011; Nickerson et al., 2018), we asked if Lgt treatment led to the accumulation of pro-Lpp, the substrate of Lgt. We initially sought to verify the various Lpp forms by leveraging a previously described protocol using SDS fractionation (Diao et al., 2017; Nakae, Ishii, & Tokunaga, 1979; Whitfield, Hancock, & Costerton, 1983). Lysozyme was added to allow for the identification of PG-linked Lpp forms, as previously demonstrated (M. Suzuki, Hara, & Matsumoto, 2002). CFT073 cell lysates were centrifuged to separate the SDS-insoluble PG-associated proteins (PAP) and SDS-soluble non-PG-associated proteins (non-PAP) (Figure 4a) and Lpp were detected by Western blot analysis. As expected, the fastest migrating form representing the triacylated mature form of Lpp (*) was enriched in the non-PAP fraction and the PG-linked Lpp forms (†) were enriched in the PAP fraction (Figure 4b). We also detected a form corresponding to the PG-linked diacylglyceryl modified pro-Lpp (DGPLP, §), as previously reported (M. Suzuki et al., 2002). We then asked if we could detect pro-Lpp in total cell lysates after Lgt depletion and used the *lspA* and *lolCDE* inducible deletion strains as controls. We confirmed that specific depletion of Lgt led to the accumulation of the unmodified pro-Lpp (UPLP, ø), (Figure 4c), consistent with previous results (Pailler, Aucher, Pires, & Buddelmeijer, 2012). While depletion of LspA led to the accumulation of DGPLP (§) and other PG-linked Lpp forms (†), depletion of LolCDE did not change the SDS-PAGE migration of Lpp as LolCDE is only critical for transport to the OM and does not affect lipoprotein biosynthesis (Figure 4c). These results now allowed us to determine whether the Lgti identified in this study inhibited Lgt in bacterial cells.

**Figure 4:**
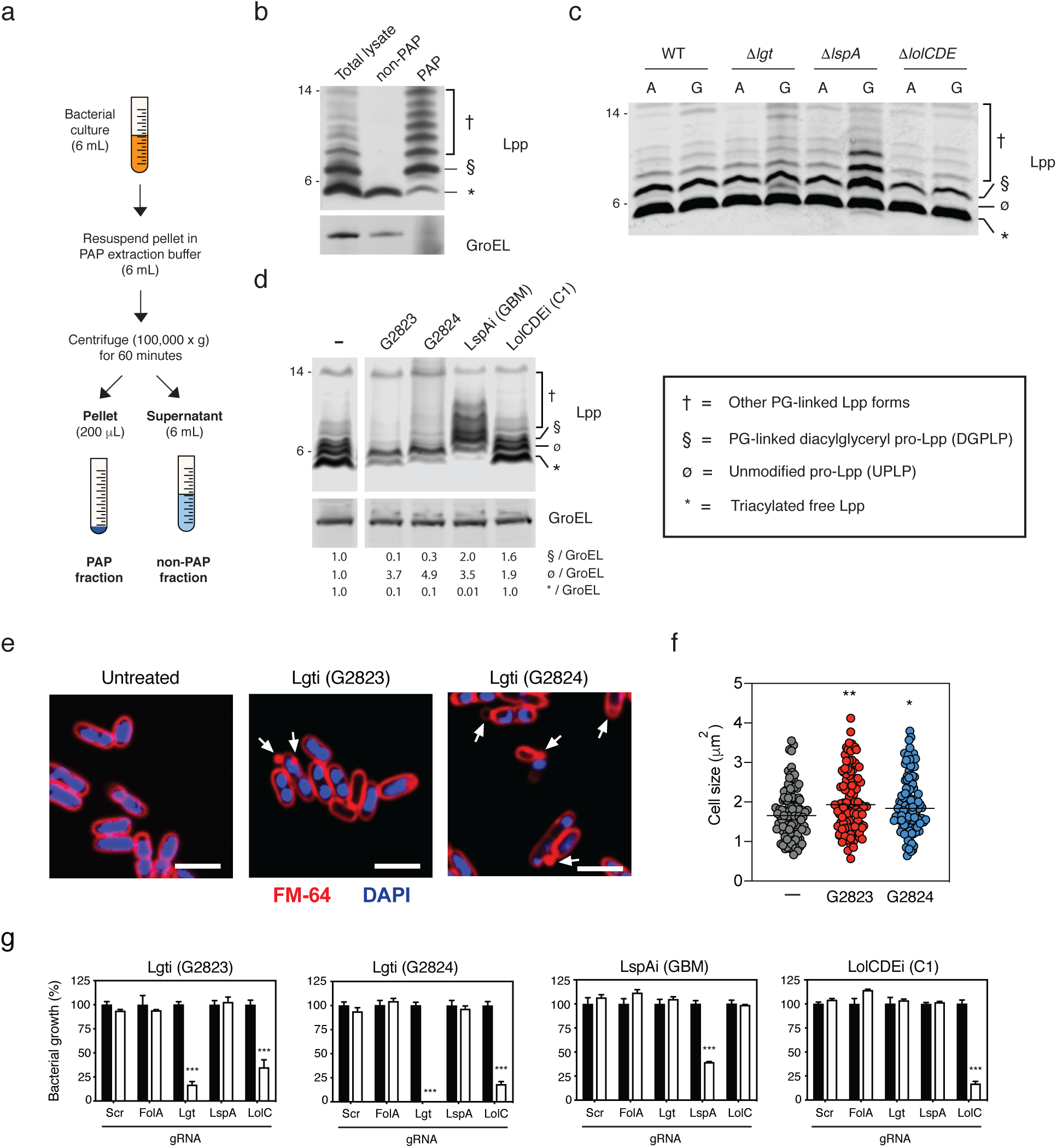
Lgti inhibit Lgt enzymatic activity in bacterial cells. **(a)** Schematic representing the isolation of PAP and non-PAP fractions. Bacterial cultures were resuspended in 6 mL of PAP extraction buffer and centrifuged at 100,000 *×* g for 60 minutes. 6 mL of supernatants were collected and pellets were resuspended in 200 μL of PAP extraction buffer. PG-linked Lpp forms were more readily detected with the concentrated PAP fractions. **(b)** SDS fractionation of WT CFT073 cells to distinguish PG-associated versus non-PG-associated forms of Lpp. CFT073*imp4213* cells were treated with SDS to enrich for PAP and non-PAP fractions as discussed in the Methods and Western blot analysis was performed to detect levels of Lpp. GroEL was used as a control for enrichment of the PAP fraction. While triacylated free Lpp (*) is enriched in the SDS-soluble non-PAP fraction, higher molecular weight Lpp species (§, †) are enriched in the SDS-insoluble PAP fraction (§ = PG-linked diacylglyceryl pro-Lpp, DGPLP; † = other PG-linked Lpp forms). Molecular weight markers (kDa) are denoted on the left of the blots. **(c)** Detection of Lpp intermediates in MG1655*Δlgt*, MG1655*ΔlspA* and MG1655*ΔlolCDE* inducible deletion strains by Western blot analysis. WT or inducible deletion strains were treated with 2% arabinose (A) or 0.2% glucose (G) and total cell lysates were harvested at 3 hours post treatment (Ø = unmodified pro-Lpp, UPLP). PG-linked DGPLP (§) and other higher molecular weight Lpp species (†) accumulated after LspA depletion. **(d)** Accumulation of pro-Lpp in cells treated with Lgti. CFT073*imp4213* cells expressing an arabinose inducible form of Lpp (*CFT073imp*4213*Δlpp:lpp*^Ara^) were incubated with arabinose for 30 minutes prior to treatment with 0.5*×*MIC concentrations of the inhibitors for another 30 minutes. Lpp forms are denoted as described above. **(e)** Lgti treatment leads to cell morphology changes and membrane blebs. CFT073*imp4213* cells were left untreated or treated with Lgti at 1×MIC for 30 minutes, fixed and incubated with FM-64 dye (red) and DAPI solution (blue) to stain membranes and nucleic acid, respectively, and visualized by confocal microscopy. Arrows represent membrane blebs and scale bars represent 3 μm. **(f)** Quantitation of cell size after treatment Lgti. A total of 104 ± 4 cells per treatment were quantitated using ImageJ (*p = 0.04; ***p = 0.002). **(g)** CRISPRi knock-down of *lgt* gene expression sensitizes cells to Lgti but not LspAi and LolCDEi. *E. coli* BW25113 cells expressing dCas9 and gRNAs specific to *lgt*, *lspA* or *lolC* were untreated (black bars) or treated (white bars) with 2 μM Lgti (G2823 and G2824), 0.05 μM LspAi (globomycin) or 0.8 μM LolCDEi (C1). A scrambled (scr) gRNA and gRNA specific to *folA* (dihydrofolate reductase) were used as negative controls. Bacterial growth was measured by OD_600_ and values were normalized to the untreated sample for each gRNA, which was set at 100% (***p < 0.001). Data are representative of at least two independent experiments each performed in triplicate.

As the Lgti have only moderate activity against WT bacterial strains, we performed mechanistic studies with Lgti in the CFT073 cells containing the *imp*4213 allele in *lptD* (CFT073*imp*4213), which leads to permeabilization of the OM (Ruiz, Falcone, Kahne, & Silhavy, 2005). Given the high expression of Lpp, we engineered CFT073*imp*4213 cells to only express an arabinose inducible *lpp* (CFT073*imp*4213*Δlpp:lpp*^Ara^) to minimize the background from pre-formed Lpp. *lpp* gene expression was induced prior to treatment with sub-MIC levels of Lgti and led to an accumulation of UPLP (ø, Figure 4d), similar to what was observed with the CFT073*Δlgt* strain (Figure 4c), and a concurrent decrease in the triacylated mature Lpp form (*, Figure 4d). While treatment with globomycin (LspAi) led to an accumulation of DGPLP and other PG-linked Lpp forms, treatment of cells with the AstraZeneca LolCDE inhibitor C1 (LolCDEi) (McLeod et al., 2015) did not lead to significant accumulation of Lpp, which is consistent with our data using the inducible deletion strains as well as published results (Narita & Tokuda, 2011; Nickerson et al., 2018). These data demonstrate that the Lgti identified in this study inhibit the generation of mature triacylated Lpp and lead to the accumulation of UPLP, which is the substrate of Lgt.

Lgt on-target activity was further confirmed using two additional methods. First, Lgti treatment also led to the expected OM blebbing and increase in cell size (Figure 4e and 4f), the former of which was previously demonstrated in a Pal-deficient *E. coli* strain (Kowata, Tochigi, Kusano, & Kojima, 2016). Second, we asked whether cells expressing reduced levels of Lgt would be specifically sensitized to Lgti compared to the other inhibitors. To test this hypothesis, we utilized CRISPRi technology to decrease gene expression of the enzymes involved in lipoprotein biosynthesis and transport. BW25113 cells containing plasmids expressing dCas9 and guide RNAs (gRNAs) specific to *lgt*, *lspA*, *lolC* were treated with Lgti, LspAi and LolCDEi and bacterial growth was measured. Scrambled (scr) and a *folA*-specific gRNAs were used as negative controls. Levels of downregulation of target gene expression (Figure 4-figure supplement 1) were consistent with published reports for CRISPRi in bacterial cells (Rousset et al., 2018). Decreased expression of *lgt* specifically sensitized cells to Lgti but not LspAi and LolCDEi (Figure 4g and Figure 4-figure supplement 2). As expected, decreased expression of *lspA* and *lolC* specifically led to enhanced growth inhibition by LspAi and LolCDEi compounds, respectively (Figure 4g and Figure 4-figure supplement 2a,b). Decreased *lolC* expression also sensitized cells to Lgti (Figure 4g) and, at higher concentrations, LspAi (Figure 4-figure supplement 2d), but we confirmed that previously identified LolCDEi-resistant mutants were not cross-resistant to Lgti (Supplemental Tble 1). Cumulatively, our data demonstrate that the novel Lgt-binding macrocycles G2823 and G2824 interfere with Lgt activity leading to inhibition of *E. coli* growth.

### Antibacterial activity of Lgti is not sensitive to *lpp* deletion

Our data with the inducible deletion strains (Figure 2h) suggested that the mechanism of cell death upon Lgt depletion is independent of Lpp (Figure 2h), distinguishing it from the mechanism of cell death after depletion of enzymes involved in later steps of lipoprotein biosynthesis. Since the Lgti identified in this study now allowed us to pharmacologically intervene at this step in the pathway, we compared the bactericidal activity of Lgti with that of LspAi and LolCDEi. We treated CFT073*imp*4213 and CFT073*imp*4213*Δlpp* cells with Lgti (G2824), LspAi (GBM) and LolCDEi (C1) at 2*×*MIC of the respective inhibitors against CFT073*imp*4213 and enumerated viable CFU counts. Consistent with our data using the inducible deletion strains, *lpp* deletion did not protect cells from Lgti (Figure 5a). In fact, the rate of CFU loss after Lgti treatment was more rapid in *lpp*-deleted cells, which is consistent with our data using the CFT073*Δlgt* cells (Figure 2h) and indicates a protective role for Lpp when targeting Lgt. As expected, inhibition of bacterial growth by LspAi and LolCDEi was lost in the absence of *lpp* (Figure 5b and 5c). In contrast, vancomycin showed equivalent killing of CFT073*imp*4213 and CFT073*imp*4213*Δlpp* at 5 hours post treatment (Figure 5d). These data confirm that *lpp* deletion is not a mechanism of resistance to Lgti, and in fact protects cells against depletion or inhibition of Lgti.

**Figure 5:**
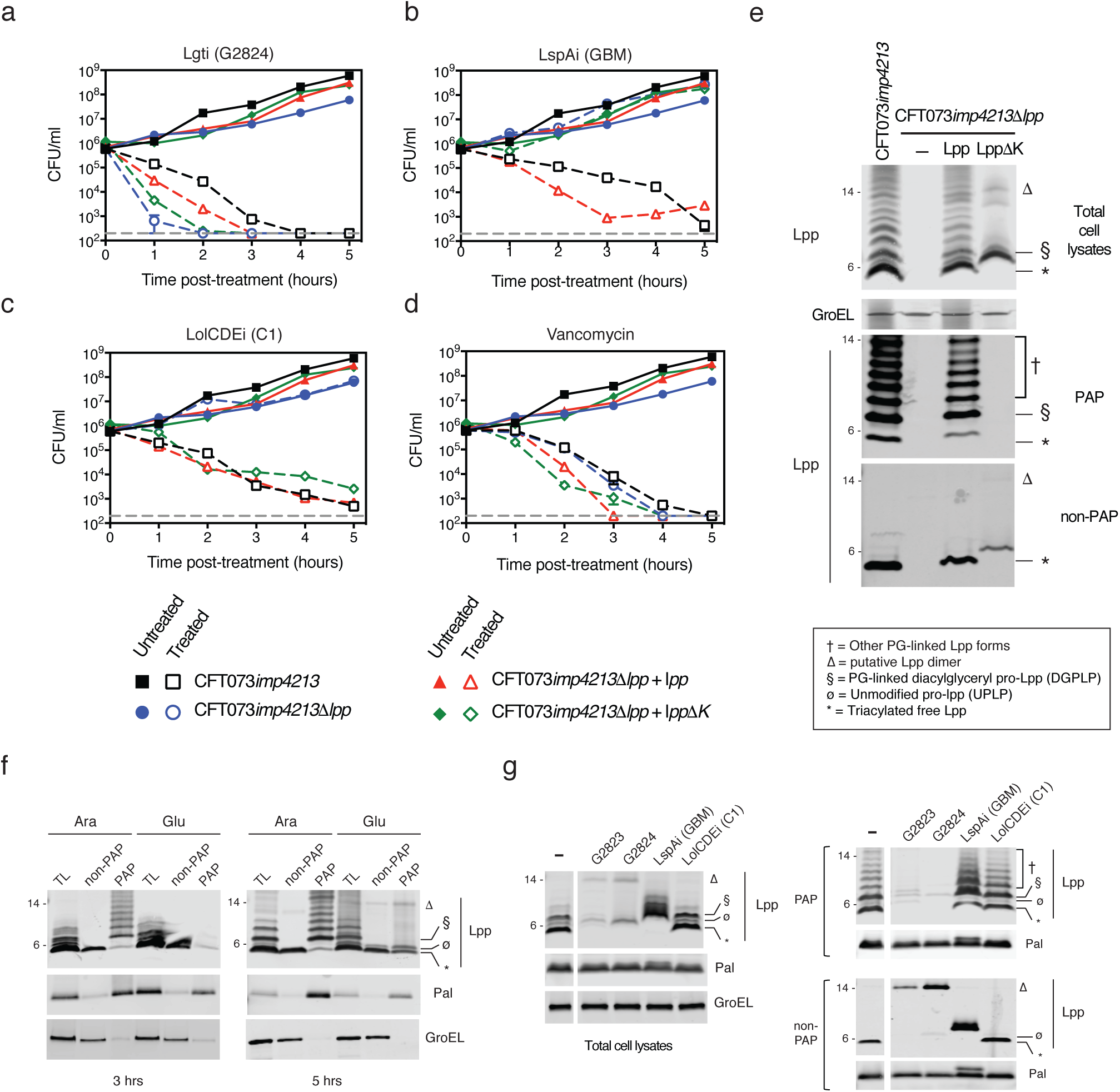
*lpp* deletion does not rescue growth after Lgti treatment. CFT073*imp*4213 (black), CFT073*imp*4213*Δlpp* (blue) or CFT073*imp*4213*Δlpp* complemented with pBAD24 plasmids encoding WT *lpp* (red) or *lppΔK* (green) were untreated (filled symbols) or treated (open symbols) with 12.5 μM Lgti G2824 **(a)**, 3.2 μM LspAi (GBM) **(b)**, 6.3 μM LolCDEi (C1) **(c)** and 1.6 μM vancomycin **(d)**. G2823 was not be tested due to limitations in compound availability. CFUs were enumerated at various times post treatment. **(e)** CFT073*imp*4213, CFT073*imp*4213*Δlpp* or CFT073*imp*4213*Δlpp* complemented with WT *lpp* or *lppΔK* were treated with SDS to enrich for PAP and non-PAP fractions. As the PAP fraction is 30-fold more concentrated than the non-PAP fraction, it is more appropriate to compare different mutants within the same fraction. Lpp forms are denoted as previously described (* = Triacylated free Lpp; § = PG-linked DGPLP; ø = UPLP; † = other PG-linked Lpp forms; *Δ* = putative Lpp dimer). The identity of the band in Figure 5e below the putative Lpp dimer (*Δ*) is unknown and could represent a degradation product. The Lpp*Δ*K is his-tagged and hence migrates slower on SDS-PAGE relative to the mature triacylated Lpp. Data are representative of three independent experiments. **(f)** Lgt depletion leads to loss of PG-linked Lpp and Pal. CFT073*Δlgt* inducible deletion cells were grown in arabinose (Ara) or glucose (Glu) and Lpp and Pal expression was determined in total cell lysates (TL), SDS-insoluble (PAP) and SDS-soluble (non-PAP) fractions at 3 and 5 hours post treatment. GroEL was used as a control for fractionation. Lpp forms are denoted by symbols as described in Figure 5e. **(g)** Lgti treatment leads to loss of PG-associated Lpp and Pal. CFT073*imp*4213 cells were treated with Lgti (G2823 and G2824), LspAi (GBM) or LolCDEi (C1) for 30 minutes at 0.5×MIC and levels of Lpp and Pal were measured in total cell lysates, PAP and non-PAP fractions. Lpp forms are denoted by symbols as described above.

To determine if PG-linkage of Lpp plays a role in protection against Lgti, we treated CFT073*imp*4213*Δlpp* cells complemented with either WT *lpp* or a mutant form that is unable to covalently link to PG (*lppΔK*). Using the previously described SDS fraction protocol, we confirmed that while WT Lpp localized to both PAP and non-PAP fractions, the Lpp*Δ*K mutant was only detected in the non-PAP fraction (Figure 5e). As we noted earlier, PG-linked DGPLP (§) and other PG-linked Lpp forms (†) were primarily detectable in cell lysates and PAP fraction (Figure 5e). While complementation of CFT073*imp*4213*Δlpp* with WT *lpp* led to increased bactericidal activity of LspAi and LolCDEi, bactericidal activity of Lgti and LolCDEi in cells deleted for *lpp* or those only expressing Lpp*Δ*K was comparable (Figure 5a-c). These data suggest that while PG-linked Lpp is toxic to cells after treatment with LspAi, it functions as a protective mechanism against Lgti. Furthermore, accumulation of PG-linked Lpp does not fully explain the bactericidal activity of LolCDEi.

### Lgt depletion or inhibition leads to decreased PG-association of Lpp and Pal

Unlike with inhibitors of LspA (Yakushi, Tajima, Matsuyama, & Tokuda, 1997a) and LolCDE (Nickerson et al., 2018), Lpp protects cells from Lgti suggesting that the PG-linkage state and/or localization of Lpp must differ after treatment with Lgti. Using SDS fractionation to enrich for PG-associated proteins in the CFT073*Δlgt* inducible deletion strain, we find that while Lgt depletion leads to a significant loss of DGPLP and other PG-linked Lpp forms in the PAP fractions, there is a modest accumulation of UPLP in the PAP fraction (Figure 5f). Lgt depletion also led to decreased PG-associated Pal, although the difference between pro-Pal and mature Pal forms was difficult to distinguish by SDS-PAGE due to larger size of Pal compared to Lpp. We then tested if Lgti treatment also led to a similar loss of PG-association of Lpp and Pal. As before, we used cells expressing an inducible form of *lpp* (CFT073*imp*4213*Δlpp:lpp*^Ara^) and find that Lgti treatment leads to decreased PG-associated DGPLP and other PG-linked Lpp forms Lpp (Figure 5g). In addition, we also detect a modest decrease in PG-associated Pal (Figure 5g). As expected, LspAi treatment led to the accumulation of PG-linked DGPLP (§) and other PG-linked Lpp forms (†). These data suggest that the accumulated UPLP after Lgt inhibition is either not significantly linked to PG or does not accumulate to levels needed to induce cell death. To address the first question, we engineered cells to only express a mutant of Lpp that has the conserved cysteine modified (CFT073*imp*4213*Δlpp*:*lpp* ^C21A^) and asked if this form was PG-linked. Lpp ^C21A^ cannot be modified by Lgt and represents the pro-Lpp substrate of Lgt. While complementation of CFT073*imp*4213*Δlpp* with WT Lpp led to normal PG-linkage, CFT073*imp*4213*Δlpp*:*lpp* ^C21A^ cells showed a significantly less PG-association of Lpp (Figure 5-figure supplement 1). Cumulatively, these data demonstrate that inhibition of Lgt leads to decreased PG-association of Lpp, which could explain why deletion of *lpp* does not lead to resistance to Lgti.

### Inhibition of Lgt does not lead to significant accumulation of PG-linked Lpp in the IM

We then asked if membrane localization of Lpp, other OM lipoproteins and OMPs were affected by Lgti. We utilized sucrose gradient centrifugation to separate *E. coli* IM and OM and measured levels of OM lipoproteins (Lpp, Pal, BamD) and OMPs (BamA, OmpA) by Western blot analyses. Sucrose gradient centrifugation of CFT073*imp*4213 cells membranes led to the efficient separation of IM and OM, as measured by MsbA and OmpA expression, respectively (Figure 6a). In comparison to untreated cells, Lgti treatment led to significant reductions of Lpp in the OM (Figure 6a). Although Lgti treatment led to accumulation of Lpp in the IM, the levels were significantly lower than that seen with LspAi and LolCDEi. All the inhibitors led to decreased OM localization of other lipoproteins, Pal and BamD, as well as the OM *β*-barrel proteins BamA and OmpA (Figure 6 and Figure 6 – figure supplement 1). These results are not totally unexpected given OmpA insertion into the OM requires BamA function, which itself requires other Bam lipoproteins, including BamD, for proper OM localization. These results suggest that while Lgti are similar to LspAi and LolCDEi in their effects on Pal and other lipoproteins involved in OM biogenesis pathways, they differ from LspAi and LolCDEi in that they do not lead to significant accumulation of Lpp in the IM supporting our data that *lpp* deletion does not play major role in resistance to Lgti.

**Figure 6:**
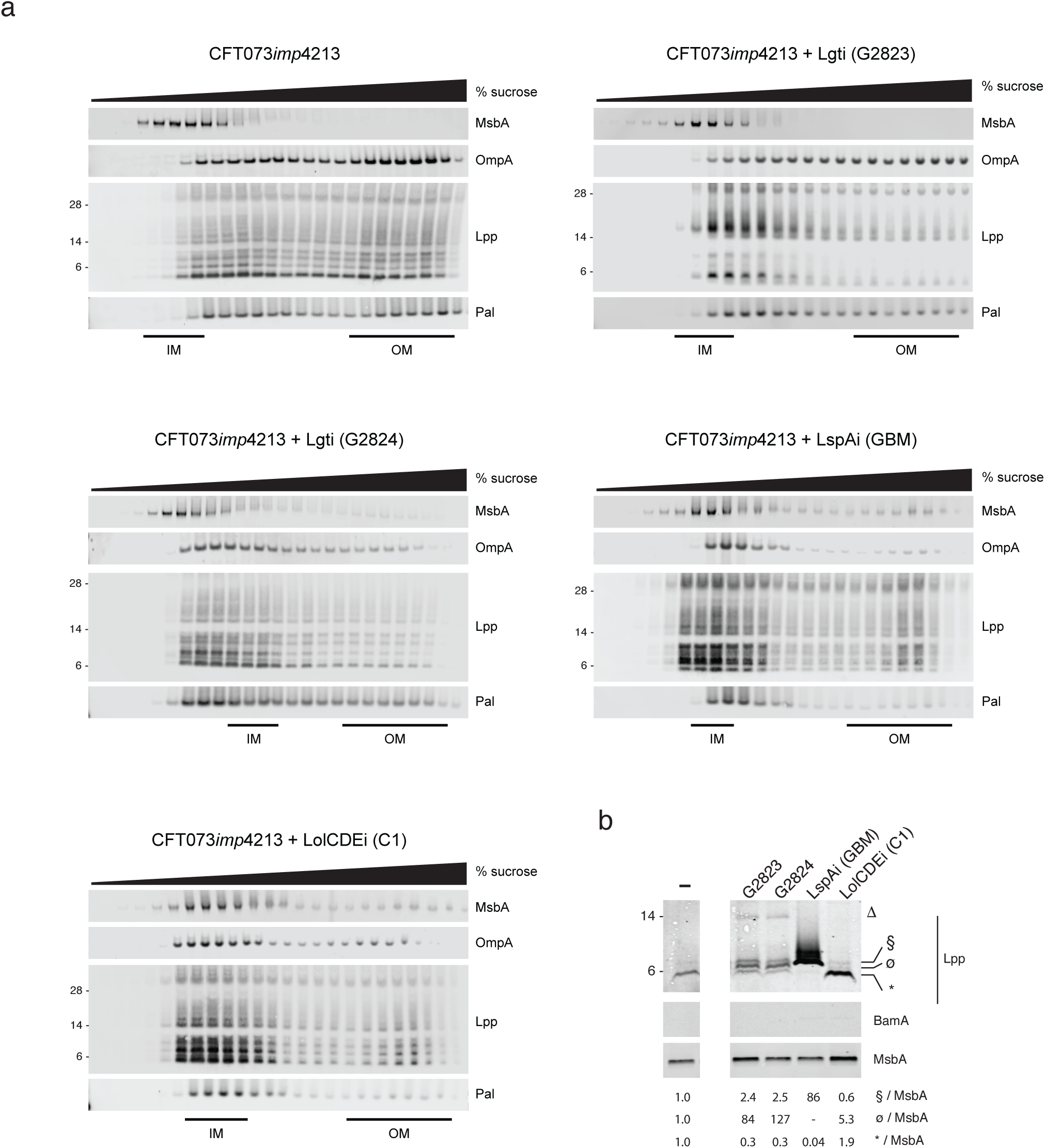
Inhibition of Lgt leads to depletion of essential OM lipoproteins and OMPs and minimal IM accumulation of PG-linked DGPLP. **(a)** CFT073*imp*4213 cells were treated with Lgti (G2823 and G2824), LspAi (GBM) or LolCDEi (C1) for 60 minutes at 1×MIC and subjected to sucrose gradient ultracentrifugation as described in the Methods. IM and OM fractions were assigned based on the expression of MsbA and OmpA, respectively. These data re representative of at least three independent experiments. **(b)** CFT073*imp*4213 cells were treated with Lgti (G2823 and G2824), LspAi (GBM) or LolCDEi (C1) and IM were solubilized using sarkosyl. Lpp and Pal levels were probed using Western blot analyses. Lpp forms denoted in the figure are as follows (* = triacylated free Lpp; § = PG-linked DGPLP; ø = UPLP; † = other PG-linked Lpp forms; *Δ* = putative Lpp dimer). IM fractions were probed for MsbA and BamA as controls. Levels of triacylated free Lpp (*), UPLP (ø) and DGPLP (§) were quantitated by normalizing to MsbA and levels detected in untreated cells (-) were set at 1.

In addition to sucrose gradient centrifugation, we also treated cells with sarkosyl that specifically solubilizes the IM and has been used for IM proteomic analyses in multiple Gram-negative bacteria (Ferrer-Navarro, Ballesté-Delpierre, Vila, & Fàbrega, 2016; Filip, Fletcher, Wulff, & Earhart, 1973; Hobb, Fields, Burns, & Thompson, 2009; Jabbour et al., 2010). Compared to untreated cells, treatment with Lgti led to a ∼84 to 127-fold increase in levels of UPLP in the IM. In contrast, DGPLP levels in the IM increased by a modest ∼2.5-fold in comparison to LspAi treatment, which results in a ∼86-fold increase of DGPLP in the IM (Figure 6b). These results confirm that Lgti treatment leads to minimal accumulation of inefficiently PG-linked UPLP in the IM, but no significant accumulation of other PG-linked Lpp forms, including the DGPLP.

## Discussion

Lipoprotein biosynthesis is a critical pathway involved in the biogenesis and maintenance of the Gram-negative bacterial OM, and disruption of any step in this pathway leads to loss of cell viability. Lpp maintains the integrity of the Gram-negative bacterial cell surface by covalent interaction between the C-terminal lysine and the *meso*-diaminopimelic acid residue of the PG layer (Braun & Wolff, 1970; Hirota, Suzuki, Nishimura, & Yasuda, 1977; H. Suzuki et al., 1978; Zhang & Wu, 1992; Zhang, Inouye, & Wu, 1992). Published data suggest that *lpp* deletion leads to rescue of growth after inhibition of LspA and LolCDE (McLeod et al., 2015; Nickerson et al., 2018; Xiao et al., 2012; Yakushi, Tajima, Matsuyama, & Tokuda, 1997a; Zwiebel et al., 1981) as well as rescue of the temperature-sensitive *Salmonella typhimurium lgt* and *lnt* mutants (Gan, Gupta, Sankaran, Schmid, & Wu, 1993; Gupta, Gan, Schmid, & Wu, 1993). While *E. coli lnt* is essential in the absence of *lpp* (Robichon et al., 2005), Lpp overexpression in the *E. coli lnt* mutant leads to bacterial cell growth arrest (Narita & Tokuda, 2011). Following up on data from Pailler et al., who demonstrated that *lgt* is essential in BW25113, a derivative of *E. coli* K-12 strain BD792, and that Lgt depletion leads to increased DNA leakage from the cell pole (Pailler et al., 2012), we demonstrate that Lgt depletion in the clinical *E. coli* strain CFT073 leads to significant perturbations to the bacterial cell envelope leading to increased sensitivity to antibiotics (Table 1), increased serum killing (Figure 2d) and attenuated virulence *in vivo* (Figure 2g).

Based on this information and published reports, we had expected that deletion of *lpp* would also lead to rescue of growth after depletion or pharmacologic inhibition of Lgt, but our data demonstrate Lpp is in fact protective in cells treated with Lgti (Figure 5) or after Lgt depletion in the CFT074*Δlgt* inducible deletion strain (Figure 2h). Both Lgt depletion and inhibition leads to the loss of PG tethering to the OM mediated by Lpp and Pal (Figure 5). Data demonstrating that cells expressing the Lpp^C21A^ mutant contain significantly less PG-linked Lpp further supports our conclusions that Lpp linkage to PG is negatively affected by Lgti. Although alternative hypotheses remain to be tested, our findings suggest that efficient crosslinking of Lpp to PG occurs only after diacylglyceryl modification of lipoprotein substrates by Lgt. Given that Lpp is critical for cell envelope stiffness (Mathelié-Guinlet, Asmar, Collet, & Dufrêne, 2020), we propose that in the absence of significant accumulation of DGPLP or other PG-linked Lpp forms, Lpp is protective against Lgti bactericidal activity. While our data is consistent with a previous report demonstrating UPLP and DGPLP are both linked to a single muropeptide unit (M. Suzuki et al., 2002), we show that the level of PG-linkage is significantly less efficient in the absence of diacylglyceryl modification of pro-Lpp. The consequence of these findings is that targeting Lgt as a novel antibacterial target would overcome a major liability of targeting other steps in the lipoprotein biosynthetic pathway, namely the off-target resistance mediated by *lpp* deletion (McLeod et al., 2015; Xiao et al., 2012; Zwiebel et al., 1981).

The Lgti identified in this study are the first described inhibitors of the first committed step in bacterial lipoprotein biosynthesis. G2823 and G2824 inhibit growth of WT *E. coli* and *A. baumannii*. We used a combination of biochemical and genetic strategies (Figure 3) to confirm these molecules function through inhibition of the diacylglyceryl transferase activity of Lgt. First, the Lgti in this study were identified using a Lgt binding screen and confirmed to inhibit Lgt enzymatic function *in vitro*. Second, the multiple effects and phenotypes detected in Lgti-treated cells were recapitulated using *lgt* inducible deletion strains, strongly arguing against off-target effects as the main cause of cell death. While we were unable to raise on-target resistant mutants to any Lgti, one could speculate that if the Lgti bind to the conserved phosphatidylglycerol binding site in Lgt, mutations disrupting Lgti binding might result in loss of Lgt function leading to cell death. This hypothesis is actually consistent with data using globomycin or an improved analog, G0790, which binds a highly conserved active site (Vogeley et al., 2016) and for which no on-target resistance mutations have ever been described (Lehman & Grabowicz, 2019; Pantua et al., 2020). As recent publications have revealed significant insights into the potential mechanisms of diacylglyceryl modification by Lgt (Mao et al., 2016; Singh et al., 2019), further studies aimed at determining if these Lgti competitively inhibit binding of the phosphatidylglycerol or prolipoprotein substrates would be critical in better understanding the mechanism by which these molecules interfere with this critical OM biogenesis pathway. Although we do not detect MIC shifts with Lgti in cells overexpressing *lgt* (data not shown), drug resistance in *E. coli* after target overexpression can increase, remain unchanged or decrease depending on the balance between bacterial fitness costs and inhibition of enzymatic activity (Palmer & Kishony, 2014). One could speculate that even a modest inhibition of Lgt could lead to significant effects on OM integrity and cellular fitness which may counteract any resistance arising from *lgt* overexpression. While CRISPRi-mediated downregulation of *lgt* expression specifically sensitizes cells to Lgti but not LspAi or LolCDEi (Figure 4g), *lolC* downregulation increases sensitivity to growth inhibition by LolCDE, Lgti and, at higher concentrations, LspAi (Figure 4-supplement 1). It is possible that LolC depletion may increase the permeability of cells to Lgti and LspAi as Lol depletion by CRISPRi has been demonstrated to lead to increased risk of plasmolysis and membrane reorganization (Caro, Place, & Mekalanos, 2019). As with many early antibiotic leads, we cannot fully rule out that the Lgti identified in this study may have additional targets in bacterial cells at higher concentrations, but our data strongly suggest that Lgti concentrations that inhibit growth of OM-permeabilized *E. coli* are consistent with their inhibition of Lgt enzymatic activity.

In addition to the fact that *lpp* deletion does not play a role in resistance to Lgti, our studies have uncovered additional novel findings that spur new questions and investigations. First, both Lgt depletion as well as pharmacologic inhibition of Lgt led to accumulation of a ∼14 kDa Lpp isoform in the IM (*Δ*, Figure 5f,g). While the identity or function of this Lpp form is unknown, its size and the fact that a Lpp form around the same molecular weight is also detected in cells expressing the Lpp*Δ*K mutant (Figure 5 – figure supplement 1) suggest that it could represent a stable Lpp dimer. While stable Lpp trimers have been described (Bjelić, Karshikoff, & Jelesarov, 2006; Shu, Liu, Ji, & Lu, 2000), there is also evidence that Lpp can exist as a dimer (Chang, Lin, Wang, & Liao, 2012). Second, while our data suggest that the lack of diacylglyceryl modification by Lgt generates a less optimal substrate for the L,D-transpeptidases that covalently link Lpp to PG, it does not rule out the possibility that PG-linkage may occur at or after modification by Lgt. Third, we demonstrate that LolCDEi remain bactericidal against cells expressing only Lpp*Δ*K. While the sucrose gradient and sarkosyl solubilization centrifugation studies demonstrate that LspAi treatment leads to accumulation of DGPLP in the IM (Figure 6), consistent with previous data with globomycin (M. Suzuki et al., 2002), no such accumulation is detected after treatment with LolCDEi. This raises the possibility that accumulation of either WT Lpp or Lpp*Δ*K could compete with less abundant essential OM lipoproteins for limited transport via LolCDE, which is consistent with our data demonstrating decreased OM localization of BamD after LolCDEi treatment (Figure 6 – figure supplement 1). Lgti, like LspAi and LolCDEi, had significant effects on OM localization of the *β*-barrel protein, OmpA (Figure 6a), which is most likely due to decreased OM expression of BamD and consequently BamA (Figure 6 – figure supplement 1). These results inform us that while Lgti behave very similarly to LspAi and LolCDEi in terms of depleting OM lipoproteins and OMPs, their effects on Lpp set them apart from other inhibitors on this pathway.

In summary, our study is the first to systematically differentiate the role of Lpp in targeting multiple steps of bacterial lipoprotein biosynthesis and transport. The loss of PG-association of Lpp and Pal, resulting from Lgt depletion or pharmacologic inhibition of Lgt, leads to significant OM defects. The identification and characterization of these Lgti validates Lgt as a novel and druggable antibacterial target and could serve as initial starting points for ongoing medicinal chemistry efforts to improve antibacterial potency and physiochemical properties. Our studies suggest that therapeutic targeting of Lgt over other steps in the lipoprotein biosynthesis and transport pathways might present a more favorable resistance profile and prevent the spread of multi-drug resistant bacterial infections.

## Materials and Methods

### Ethics statement

All mice used in this study were housed and maintained at Genentech in accordance with American Association of Laboratory Animal Care guidelines. All experimental studies were conducted under protocol 13-0979A were approved by the Institutional Animal Care and Use Committee of Genentech Lab Animal Research and performed in an Association for Assessment and Accreditation of Laboratory Animal Care International (AAALAC)-accredited facility in accordance with the Guide for the Care and Use of Laboratory Animals and applicable laws and regulations.

### Antibodies

The anti-Pal antibody was a generous gift from Dr. Shaw Warren (Massachusetts General Hospital). The anti-OmpA (Antibody Research Corporation), anti-GroEL (Enzo Life Sciences), anti-ThyA (GeneTex, Inc) and anti-His (Cell Signaling Technology) antibodies were obtained from commercial sources. Generation of anti-Lpp and anti-BamA antibodies has been previously described (Diao et al., 2017; Storek et al., 2018; 2019). Recombinant Lgt and BamD were used to generate rabbit polyclonal antibodies. Rabbit immunizations, generation of antisera and purification of rabbit polyclonal antibodies were performed as previously described for Lpp (Diao et al., 2017).

### Generation of bacterial strains and plasmids

Bacterial strains and plasmids used in this study are listed in Supplemental File 2. *E. coli* strain CFT073 (ATCC 700928) (Mobley et al., 1990) and MG1655 (ATCC 700926) were purchased from ATCC. Gene disruption in CFT073 was performed as previously described (Datsenko & Wanner, 2000; Diao et al., 2017). CFT073*Δlgt* was generated based on the previously published protocol (Pailler et al., 2012) by retaining the *lgt* stop codon, which forms part of the *thyA* ribosomal binding site. The primers used to generate the CFT073 and MG1655 mutants are listed in Supplemental File 3. Plasmids pKD46 for the *l* Red recombinase (Datsenko & Wanner, 2000; Diao et al., 2017), pKD4 or pSim18 for the integration construction (Datsenko & Wanner, 2000; Diao et al., 2017) and pCP20 (Cherepanov & Wackernagel, 1995) for the FLP recombinase were used in this study. The inducible deletion strains (MG1655*Δlgt,* MG1655*ΔlspA* and MG1655*ΔlolCDE*) in either the WT and/or *Δlpp* backgrounds were generated using similar methods as previously described (Diao et al., 2017; Noland et al., 2017). CFT073*imp*4213*Δlpp* containing pBAD24-*lpp* was used to generate CFT073*imp*4213*Δlpp:lpp^Ara^* to detect the different Lpp species after treatment with pharmacologic inhibitors. The PA14*imp*4213 strain was generated based on published protocols (Balibar & Grabowicz, 2016; Hmelo et al., 2015). For expression under the IPTG-inducible promoter, DNA encoding the full-length sequences of *lgt*, *lspA*, *lnt* and *lolCDE* were cloned into pLMG18 and induced using 2.5 mM IPTG.

### *In vitro* growth inhibition and serum sensitivity assays

Unless stated otherwise, *E. coli* cells were grown in Luria-Bertani (LB) medium (0.5% yeast extract, 1% tryptone, 0.5% NaCl) at 37°C. When indicated, kanamycin (Kan) was added to culture media at a 50 μg/ml final concentration. MIC assays were performed based on Clinical and Laboratory Standards Institute (CLSI) guidelines. For *in vitro* growth curves, overnight cultures of WT CFT073, CFT073*Δlgt* and CFT073*Δlgt* complemented with *lgt* from *E. coli* (*lgt* ^Ec^) or *P. aeruginosa* (*lgt* ^Pa^) were grown to mid-exponential phase (OD_600_=0.6) and then diluted to OD_600_=0.1 to initiate growth curves. At various times, culture aliquots were diluted and plated in dilutions on LB+Kan agar and CFUs were enumerated in duplicate. Growth of MG1655 inducible deletion strains was measured by culturing in the presence of two-fold dilutions of arabinose (starting arabinose concentrations for CFT073 and MG1655 inducible deletion strains were 4% and 0.8%, respectively). While 2% arabinose was sufficient for WT growth of MG1655*Δlgt,* 4% arabinose was used for CFT073*Δlgt* based on comparing its growth to that of WT CFT073 as measured by CFUs. OD_600_ growth measurements were performed using an EnVision 2101 Multilabel Reader plate reader (PerkinElmer) linked with Echo Liquid Handler (Labcyte). For time-kill experiments, bacteria were harvested in mid-exponential phase and treated with 12.5 μM Lgti G2824, 3.2 μM LspAi (GBM), 6.3 μM LolCDEi (C1) and 1.6 μM vancomycin. G2823 was not be tested due to limitations in compound availability. CFUs were enumerated at various times post treatment. Bacterial culture medium containing 2% arabinose was used to induce *lpp* or *lppΔ*K expression from pBad24 plasmids. Bacterial viability at different time points during the treatment was measured by enumerating CFU. Serum killing assays were carried out as previously described (Diao et al., 2017).

### Detection of membrane permeability using SYTOX Green incorporation

To determine the effect of Lgt depletion on membrane permeability, WT CFT073 and CFT073*Δlgt* strains were streaked onto a LB agar plate containing 4% arabinose and cultured at 37°C for 18 hours. From a single colony, bacteria were cultured in LB broth containing 4% arabinose and cultured at 37°C to OD 0.5. One mL cultures of OD=0.5 for both strains were harvested, washed and resuspended in LB broth or medium containing a range of arabinose (2, 0.2, 0.1, 0.05% and 0) or glucose (0.2%) concentrations and incubated at 37°C for 2 hours. Cells were harvested by centrifugation at 4000 *×* g at 4°C for 5 minutes. Intact CFT073 or CFT073 treated with 70% ethanol at RT for 15 minutes to permeabilize the cells were used as controls. Cells were incubated with SYTOX green following the manufacturer’s recommendation, washed with PBS (3*×*) and fixed in 2% paraformaldehyde. SYTOX Green incorporation was measured by flow cytometry using a FACS Aria II (Becton Dickenson) and analyzed using Flowjo software.

### Mouse infection model

Overnight bacterial cultures were back diluted 1:100 in M9 media and grown to an OD_600_=0.8-1 at 37°C. Cells were harvested, washed once with PBS and resuspended in PBS containing 10% glycerol. Cells were frozen in aliquots and thawed aliquots were measured for CFUs prior to mouse infections. Virulence of WT CFT073 and CFT073*Δlgt* was measured using the neutropenic *E. coli* infection model (Cross, Siegel, Byrne, Trautmann, & Finbloom, 1989). Seven-week-old female A/J mice (Jackson Laboratory) were rendered neutropenic by peritoneal injection of 2 doses of cyclophosphamide (150 mg/kg on Day -4 and 100 mg/kg on Day -1). On Day 0, mice were infected with 5 *×*10^5^ CFU of mid-exponential phase bacteria diluted in PBS by intravenous injection through the tail vein. At 30 minutes and 24 hours post infection, bacterial burden in the liver and spleen was determined by serial dilutions of tissue homogenates on LB plates.

### Macrocyclic peptide library design and selection of Lgt-binding molecules

A thioether-macrocyclic peptide library was constructed by using N-chroloacetyl D-phenylalanine (ClAc-f) as an initiator in a genetically reprogrammed in vitro translation system (Kashiwagi et al., 2013). The genetic code was designed with the addition of two N-methyl amino acids: N-methyl-L-phenylalanine (MeF) and N-methyl-L-glycine (MeG); and, three unnatural amino acids: (S)-2-aminoheptanoic acid (Ahp), (S)-3-([1,1’-biphenyl]-4-yl)-2-aminopropanoic acid (Bph), and (S)- 1,2,3,4-tetrahydroisoquinoline-3-carboxylic acid (Tic) in addition to 11 natural amino acids (Ser, Tyr, Trp, Leu, Pro, His, Arg, Asn, Val, Asp, and Gly). After *in vitro* translation, a thioether bond formed spontaneously between the N-terminal ClAc group of the initiator D-phenylalanine residue and the sulfhydryl group of a downstream cysteine residue to generate the macrocyclic peptides.

Affinity selection of macrocyclic peptides binding to Lgt was performed using *E. coli* Lgt-biotin in 0.02% n-dodecyl β-D-maltoside (DDM). Briefly, 10 μM mRNA library was hybridized with a peptide-linker (11 μM) at RT for 3 minutes. The mRNA library was translated at 37°C for 30 minutes in the reprogrammed *in vitro* translation system to generate the peptide-mRNA fusion library (Goto et al., 2011; Ishizawa et al., 2013). Each reaction contained 2 μM mRNA-peptide-linker conjugate, 12.5 μM initiator tRNA (tRNAfMet aminoacylated with ClAc-D-Phe), and 25 μM of each elongator tRNA aminoacylated with the specified non-canonical /canonical amino acids. In the first round of selections, translation was performed at 20 μL scale. After the translation, the reaction was quenched with 17 mM EDTA. The product was subsequently reverse-transcribed using RNase H minus reverse transcriptase (Promega) at 42°C for 30 minutes and buffer was exchanged for DDM buffer: 50 mM Tris (pH 8), 5 mM EDTA, 200 mM NaCl2, 0.02% DDM, and 1 mM Glutathione. For affinity selection, the peptide-mRNA/cDNA solution was incubated with 250 nM biotinylated *E. coli* Lgt for 60 minutes at 4°C and the streptavidin-coated beads (Dynabeads M-280 Streptavidin, Thermo) were further added and incubated for 10 minutes to isolate Lgt binders. The beads were washed once with cold DDM buffer, the cDNA was eluted from the beads by heating for 5 minutes at 95°C, and fractional recovery from the affinity selection step were assessed by quantitative PCR using Sybr Green I on a LightCycler thermal cycler (Roche). After five rounds of affinity maturation, two additional rounds of off-rate selections were performed by increasing the wash stringency before elation to identify high affinity binders. Sequencing of the final enriched cDNA was carried out using a MiSeq next generation sequencer (Illumina).

### Peptide Synthesis

Thioether macrocyclic peptides were synthesized using standard Fmoc solid phase peptide synthesis (SPPS). Following coupling of all amino acids, the deprotected N-terminus was chloroacetylated on-resin followed by global deprotection using a trifluoroacetic acid (TFA) deprotection cocktail. The peptides were then precipitated from the deprotection solution by adding over 10-fold excess diethyl ether. Crude peptide pellets were then dissolved and re-pelleted 3 times using diethyl ether. After the final wash, the pellet was left to dry and then the pellet was resuspended in DMSO followed by the addition of triethylamine for intramolecular cyclization via formation of a thioether bond between the thiol of the cysteine and N-terminal chloroacetyl group. Upon completion of cyclization, the reaction was quenched with AcOH and the cyclic peptide was purified using standard reverse-phase HPLC methods.

### SDS-PAGE and Western immunoblotting

Bacterial cell samples normalized for equivalent OD_600_ and resuspended in Bugbuster lysis buffer (Fischer Scientific) with the addition of sample buffer (LI-COR), and separated by SDS-PAGE using 16% Tricine protein gels or NuPAGE 4-20% Bis-Tris gels (Thermofisher Scientific) and transferred to nitrocellulose membranes using the iBlot™2 Dry Blotting system (Invitrogen) and blocked using LI-COR blocking buffer for 30 minutes. Unless stated otherwise, loading buffer with reducing agents were added and samples were not boiled prior to SDS-PAGE. For the sucrose gradient centrifugation, samples were boiled prior to running the SDS-PAGE. Primary antibodies were used at a final concentration of 1μg/ml with some exceptions: rabbit anti-Lpp polyclonal antibody (0.1μg/ml); murine anti-Pal 6D7 (0.5 μg/ml); rabbit anti-GroEL (1:10,000 final dilution); rabbit anti-OmpA (1:50,000 final dilution). The secondary antibodies were all obtained from LI-COR, and used as per manufacturer’s instructions. Images were collected using the Odyssey CLx imaging system (LI-COR) and analyzed by Image Studio Lite.

### Expression and purification of recombinant Lgt and BamD

DNA encoding full-length *E. coli* Lgt fused to a C-terminal Flag-tag was transformed into Rosetta 2(DE3) Gold cells (Agilent). Starter cultures were grown in Terrific Broth (TB) media with carbenicillin (50μg/mL) and chloramphenicol (12.5μg/mL) at 37°C for 3 hours. The starter cultures were diluted 1:50 in TB medium with carbenicillin (50μg/mL), chloramphenicol (12.5μg/mL), and glycerol (1%) and grown at 37°C for 2 hours with shaking at 200 rpm. The temperature of the culture was reduced to 30°C and grown for an additional 2 hours before the temperature of the culture was reduced to 16°C and grown for 64 hours. The cells were harvested by centrifugation and resuspended into lysis buffer (20mM Tris, pH 8.0, 300mM NaCl, Protease Inhibitor cocktail and Lysonase) and stirred at 4°C for 30 minutes before being passed through a microfluidizer 3 times. The membrane fraction was solubilized by adding DDM directly to the lysate to a final concentration of 1% and stirring at 4°C for 2 hours before centrifugation at 40,000 rpm for 1 hour. Pre-equilibrated FLAG resin was added to the supernatant and incubated with rotation at 4°C for 2 hours. The slurry was added to a gravity column and the column was washed with 10 CV buffer A (20mM Tris, pH 8.0, 300mM NaCl, 5% glycerol, 1% DDM) and 10 CV buffer B (20mM Tris, pH 8.0, 300mM NaCl, 5% glycerol, 0.05% DDM). The bound fraction was eluted by the addition of 5 CV of buffer C (20mM Tris, pH 8.0, 300mM NaCl, 5% glycerol, 0.05% DDM, 100μg/mL FLAG peptide). The peak fractions were collected, concentrated to less than 5 mL and loaded onto a superdex 200 16/60 column equilibrated with buffer C (20mM Tris, pH 8.0, 300mM NaCl, 5% glycerol, 0.05% DDM, 1mM TCEP). The peak fractions were collected, analyzed by SDS-PAGE and stored at -80°C.

For recombinant *E. coli* BamD protein expression, DNA fragments encoding BamD (Gly_22_-Thr_245_) were cloned into a modified pET-52b expression vector containing an C-terminal His_8_-tag and overexpressed in *E. coli* host Rosetta 2 (DE3) grown by fermentation at 17°C for 64 hours, at which point cells were collected and resuspended in 50 mM Tris, pH 8.0, 300 mM NaCl, 0.5 mM TCEP containing cOmplete Protease Inhibitors (Roche), 1 mM PMSF and 2 U/ml of Benzonase nuclease (Sigma Aldrich). After cell lysis by microfluidization and low speed centrifugation, soluble protein was purified by Ni-NTA affinity and size exclusion chromatography. The peak fractions containing BamD were pooled and concentrated to 5 mg/mL.

### Development of the Lgt biochemical assay

The Lgt enzymatic activity was measured by specific detection of G3P. Both G3P and G1P are released from phosphatidylglycerol as Lgt catalyzes the transfer of diacylglyceryl from phosphatidylglycerol to the preprolipoprotein substrate, since the PG substrate used in the assay contains a racemic glycerol moiety at the end of phosphatidyl group. The standard assay consists of 6 μL reaction mixture with 3 nM Lgt-DDM, 50 μM phosphatidylglycerol (1,2-dipalmitoyl-*sn*-glycero-3-phospho-(1’-*rac*-glycerol), Avanti), 12.5 μM Pal-IAAC peptide substrate derived from the Pal lipoprotein (MQLNKVLKGLMIALPVMAIAA<u>C</u>SSNKN, synthesized by CPC Scientific) in 50 mM Tris, pH 8, 200 mM NaCl, 5 mM EDTA, 0.02% DDM, 0.05 % Bovine Skin Gelatin, and 1 mM glutathione. As a control, we used a mutant non-modifiable Pal substrate peptide containing a cysteine to alanine mutation (Pal-IAA**<u>A</u>**) which served as a competitive non-modifiable inhibitor. The reaction was quenched after 60 minutes at RT with 0.5 μL of 4.8% Lauryl Dimethylamine-N-Oxide (Anatrace), followed by addition of 6 μL Detection Solution. After incubation for 120 minutes at RT, the luminescence signal was read. The Detection Solution was modified based on a NAD Glo protocol (Promega, G9072), per manufacturer’s instruction. Specifically, 10 mL Detection Solution consists of 3-fold dilution of Luciferin Detection Reagents, supplemented with 10 μL Reductase, 2.5 μL Reductase Substrate, 1 mM NAD, and 4.25 U of G3PDH (Roche Diagnostics, 10127779001). The Luciferin Detection Reagents, Reductase, and Reductase Substrate were all from the NAD Glo kit (Promega). Luminescence values were normalized to DMSO controls (0% inhibition) and no enzyme controls (100% inhibition). IC50 values were calculated using a 4 parameter logistic model using GraphPad Prism software.

### Visualization of WT CFT073 and CFT073Δ*lgt* by time-lapse microcopy, confocal microcopy, transmission electron microscopy

Electron microscopy was performed as previously described (Noland et al., 2017). For time-lapse microscopy, WT CFT073, CFT073*Δlgt* and CFT073*ΔlgtΔlpp* cells were grown overnight in LB medium containing 4% arabinose, back-diluted to a final OD_600_ of 0.1 and immediately placed between a cover slip and 1% agarose pad containing 0.2% glucose for imaging. Cells were maintained at 37 °C during imaging in a stage top chamber (Okolab Inc.). Cells were imaged on a Nikon Eclipse Ti inverted confocal microscope (Nikon Instruments Inc.) coupled with a UltraVIEW VoX (PerkinElmer Inc.) and a 100× (NA 1.40) oil-immersion objective. Images were captured at various times using ORCA-Flash 4.0 CMOS camera (Hamamatsu Photonics), collected using Volocity software (Quorum Technologies) and processed using Fiji (Schindelin et al., 2012). For confocal microscopy, images were acquired on a Leica SP8 STED 3x platform using a 100× white light, NA:1.4 oil immersion objective. CFT073*imp4213* cells were treated with Lgti, LspAi or LolCDEi at 1×MIC for 30 minutes, fixed with 4% paraformaldehyde and incubated with 1 μg/mL FM-64 dye and 1 μg/mL DAPI solution. Quantitation of bacterial cell area was performed using the ImageJ program by measuring at least ∼100 bacterial cells from two independent experiments.

### Targeted downregulation of gene expression by CRISPRi

The two-plasmid bacterial CRISPRi system pdCas9-bacteria_GNE and pgRNA-bacteria_GNE are based off the AddGene plasmids 44249 and 44251 (Qi et al., 2013), respectively. The plasmid was synthesized in smaller DNA fragments (500bp-3kb) (IDT gBlocks) and assembled by Gibson Assembly (NEB) according to manufacturer’s protocols. Plasmids were confirmed by sequencing (ELIM Bio). gRNAs were designed to target the 5’ end of the gene on the non-template strand using Benchling CRISPR software (Peters et al., 2016). gRNAs were cloned into pgRNA-bacteria using Gibson Assembly (NEB) according to manufacturer’s protocols and sequence confirmed (ELIM Bio).

Bacterial cultures were grown overnight on LB agar supplemented with carbenicillin (50 μg/mL) and chloramphenicol (12.5 μg/mL) to maintain both plasmids, pdCas9-bacteria and pgRNA-bacteria with each gRNA as appropriate. Cells were scraped from the plate into fresh media. OD_600_ was measured and subsequently diluted to OD_600_=0.001 in the presence or absence of Lgti, LspAi and LolCDEi. 200 μL was transferred to a 96-well plate (Corning) and monitored for growth by measuring OD_600_ (EnVision Multimode Plate Reader, PerkinElmer). All treatments were performed in triplicate. Specificity of CRISPRi downregulation was measured using RT-qPCR.

### Purification of peptidoglycan-associated proteins

Purification of PG-associated proteins (PAP) was performed according to published methods (Diao et al., 2017; Nakae et al., 1979; Whitfield et al., 1983) with some modifications. Briefly, bacteria were harvested in mid-exponential phase for treatment and then subjected for PAP extraction by resuspended cell pellets from 10 OD (A_600_) in 6 mL of PAP extraction buffer containing 2% (wt/vol) SDS in 100 mM Tris-HCl (pH 8.0) with 100mM NaCl, 10% glycerol, and cOmplete^TM^, mini, EDTA-free protease inhibitor cocktail (Sigma-Aldrich). After 60 minutes at RT, the extraction was subjected to centrifugation at 100,000 *×* g for 60 minutes at 22°C, and the pellet, containing PG and associated proteins, was washed once with the same PAP extraction buffer with centrifugation at 100,000 *×*g for 30 minutes and resuspended in 200 μL of PAP extraction buffer (referred to as the SDS-insoluble on PAP fraction). The supernatant containing the SDS-soluble fraction was aliquoted and frozen (referred to as non-PAP fraction). Both fractions were treated with equal volume of BugBuster buffer prior to the addition of sample buffer for Western immunoblotting as described above. It should be noted that the final PAP fractions are ∼30-fold more concentrated than the non-PAP fractions.

### Isolation of *E. coli* IM and OM using sucrose gradient centrifugation and sarkosyl fractionation

Bacterial inner and outer membranes were separated by sucrose gradient as previously (Nickerson et al., 2018; Yakushi, Tajima, Matsuyama, & Tokuda, 1997b) with some modifications. Briefly, bacteria were grown in Luria broth at 37°C to mid-exponential phase (OD_600_=0.6), and then treated with 1*×*MIC of indicated inhibitors for 1 hour. Cells representing 30-40 OD_600_ equivalents were harvested by centrifugation at 4000 *×*g for 15 minutes, washed once with 50 mM Tris-HCl (pH 7.5) containing 25% (wt/vol) sucrose and Complete EDTA-free protease inhibitor cocktail (Roche), and then incubated for 10 minutes at RT in the same buffer containing 100 μg/ml lysozyme (Thermo Scientific) and 1000 U/ml nuclease (BenzonaseNuclease, EMD Millipore). Two-fold volume of ice-cold EDTA (pH 8.0) was added and the suspension was disrupted by two passages through an LV1 Microfluidizer (Microfluidics). Unbroken cells were removed by centrifugation at 4,000 × *g* and membranes were collected by ultracentrifugation at 100,000 × *g* for 1 hour and washed once with 50 mM Tris-HCl (pH 7.5). The final membrane preparation was resuspended in 50 mM Tris-HCl (pH 7.5) containing 10% sucrose, 1.5 mM EDTA and protease inhibitor cocktail and then applied to a 30 to 70% (wt/vol) sucrose gradient. The loaded gradients were spun at 200,000 *×* g for 22 hours at 4°C in a Beckman SW41Ti rotor. Fractions were removed and analyzed by SDS-PAGE and immunoblotting with appropriate antibodies. The IM fractionation of bacterial cells using sarkosyl was performed according to published methods (Filip et al., 1973; Pantua et al., 2020)..

### Statistical analyses

All statistical analyses were performed using GraphPad Prism 6.0 software (GraphPad). The data was tested for being parametric and statistical analyses were performed on log-transformed data. All graphs represent the mean ± the standard error of the mean (SEM). Unless stated otherwise, p values for all data were determined using regular unpaired *t* test (* = p<0.05, ** = p<0.01, and *** = p<0.001). p values for mouse CFU studies were determined using the Mann Whitney Test. Bonferroni correction was applied to control for multiple comparisons for CRIPSRi data in Figure 4g.

## Acknowledgements

We thank Dr. Shaw Warren (Massachusetts General Hospital) for the anti-Pal antibody, Erin Dueber for help with purification of BamD, Scott Stawicki for assistance with generating the rabbit polyclonal antibodies, and Eric Brown for helpful comments and suggestions for the manuscript.

## Figure supplement legends

**Figure 2-figure supplement 1:**
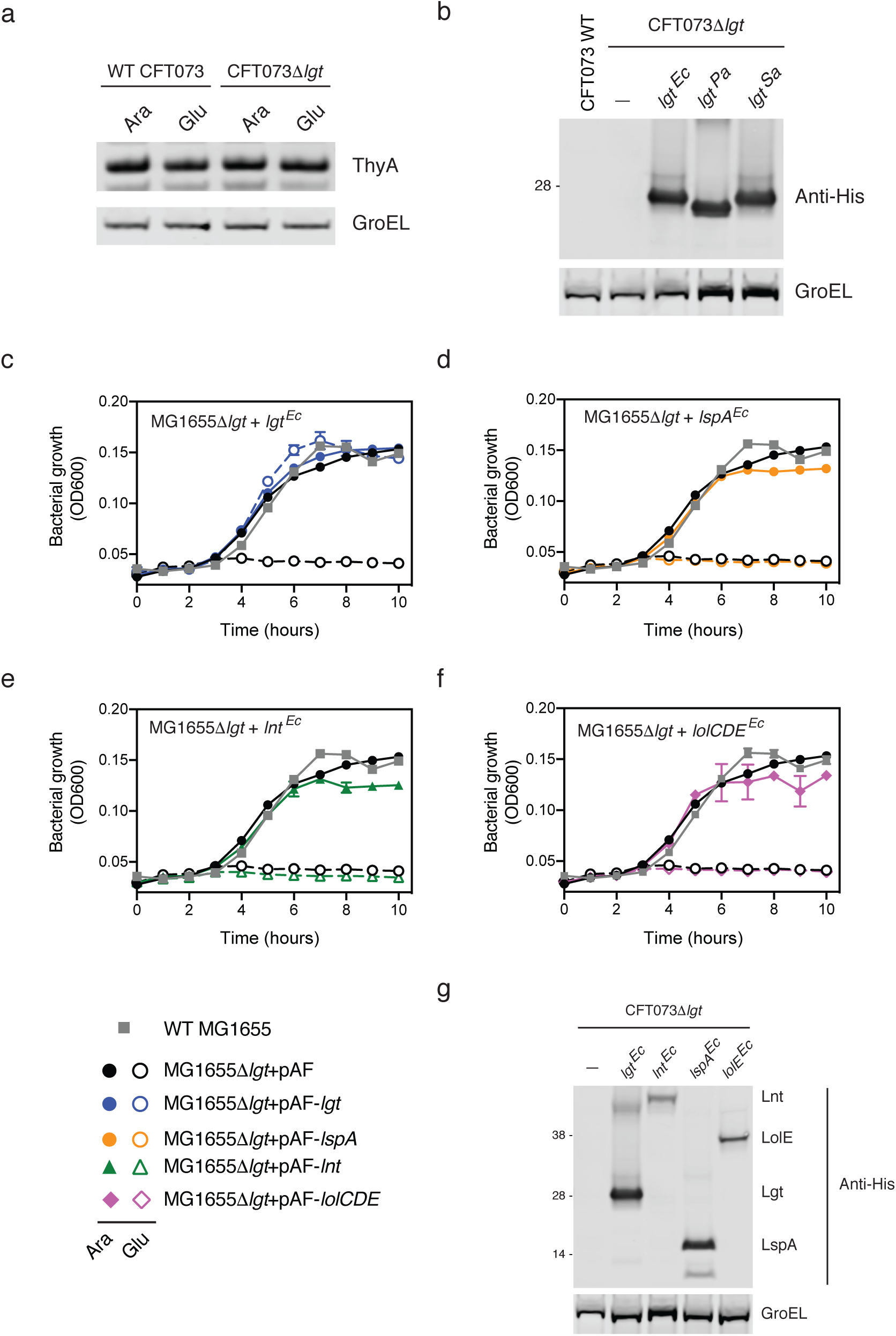
**(a)** Normal expression of thymidylate synthase (ThyA) after depletion of Lgt. WT CFT073 and CFT073*Δlgt* were grown under wild-type (4% arabinose, Ara) or depleted (0.2% glucose, Glu) conditions for 4 hours and total cell lysates were subjected to Western blot analyses using an anti-ThyA antibody. GroEL was used as a loading control. **(b)** Western blot analyses confirming protein expression after complementation with pLMG18 expressing *lgt* from *E. coli* (*lgt*^Ec^), *P. aeruginosa* (*lgt*^Pa^) or *S. aureus* (*lgt*^Sa^). All complemented *lpp* contain a c-terminal His-tag. **(c-f)** Loss of *E. coli* MG1655*Δlgt* viability after Lgt depletion is rescued after complementing with *E. coli lgt* (*lgt*^Ec^) but not *E. coli lspA* (*lspA*^Ec^), *lnt* (*lnt*^Ec^) or *lolCDE* (*lolCDE*^Ec^). 2.5 mM IPTG was used to induce expression of *E. coli lspA*, *lnt* or *lolCDE*. Cells were grown in arabinose (filled symbols, Ara) or glucose (open symbols, Glu) and bacterial growth was measured by OD_600_. **(g)** Anti-His Western blot analyses demonstrating protein expression of *E. coli* Lgt, LspA, Lnt and LolE in CFT073*Δlgt* cells complemented with His-tagged versions of the respective genes.

**Figure 2-figure supplement 2:**
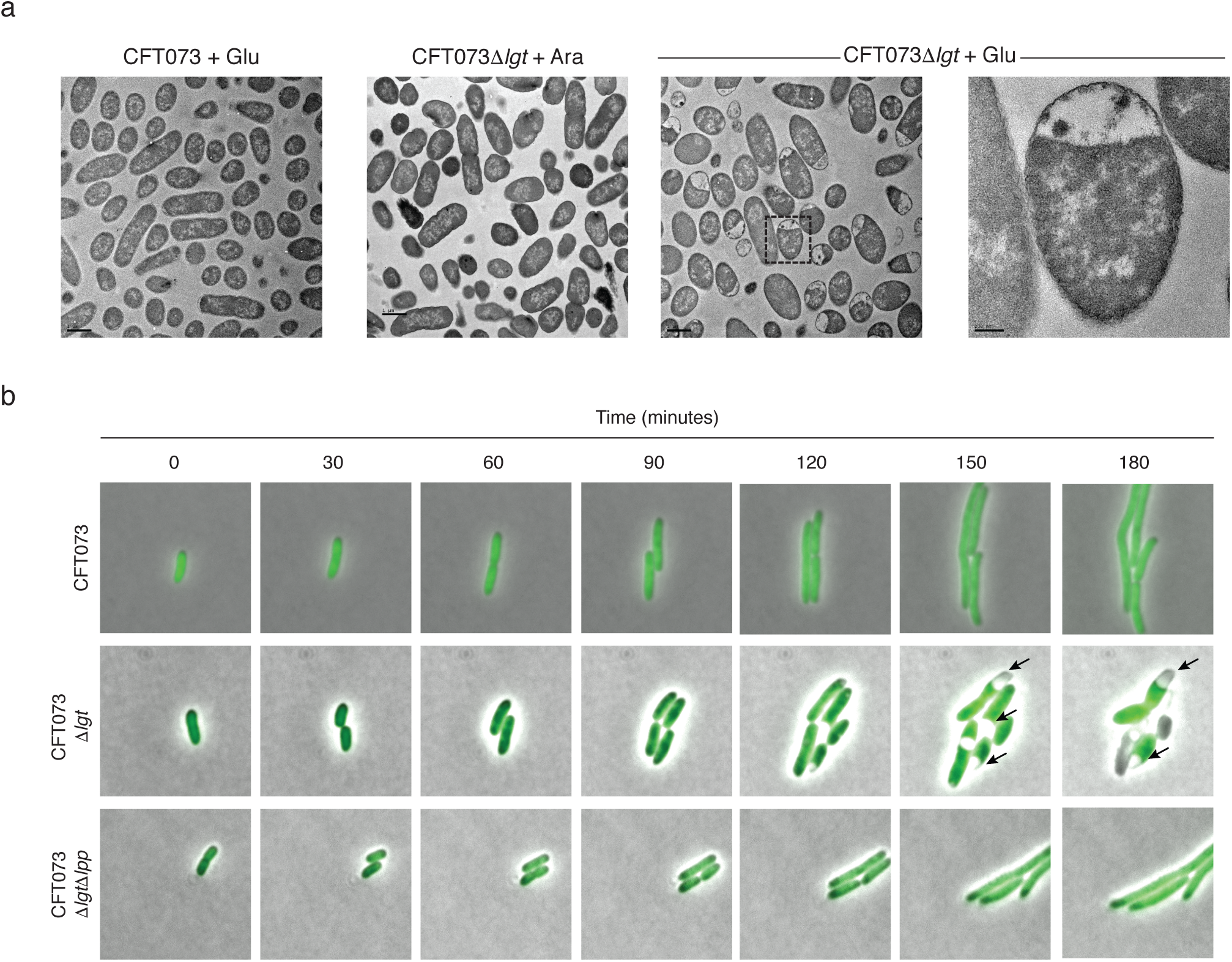
Lgt depletion results in IM contraction and the expected globular cellular phenotype. **(a)** CFT073 and CFT073*Δlgt* deletion strains were treated for 2 hours with 4% arabinose (Ara) or 0.2% glucose (Glu) and samples were processed for imaging by Transmission electron microscopy. Bars represent 1 µm (200 nm for last panel). **(b)** Live cell imaging of WT CFT073, CFT073*Δlgt* and CFT073*ΔlgtΔlpp* inducible deletion strains containing a plasmid expressing *gfp* (pGFP) were grown in the presence of 0.2% glucose. Phase contrast and fluorescence microscopy images were overlayed at various times post treatment. Arrows denote IM contraction which is not observed in the strain containing the *lpp* deletion.

**Figure 3-figure supplement 1:**
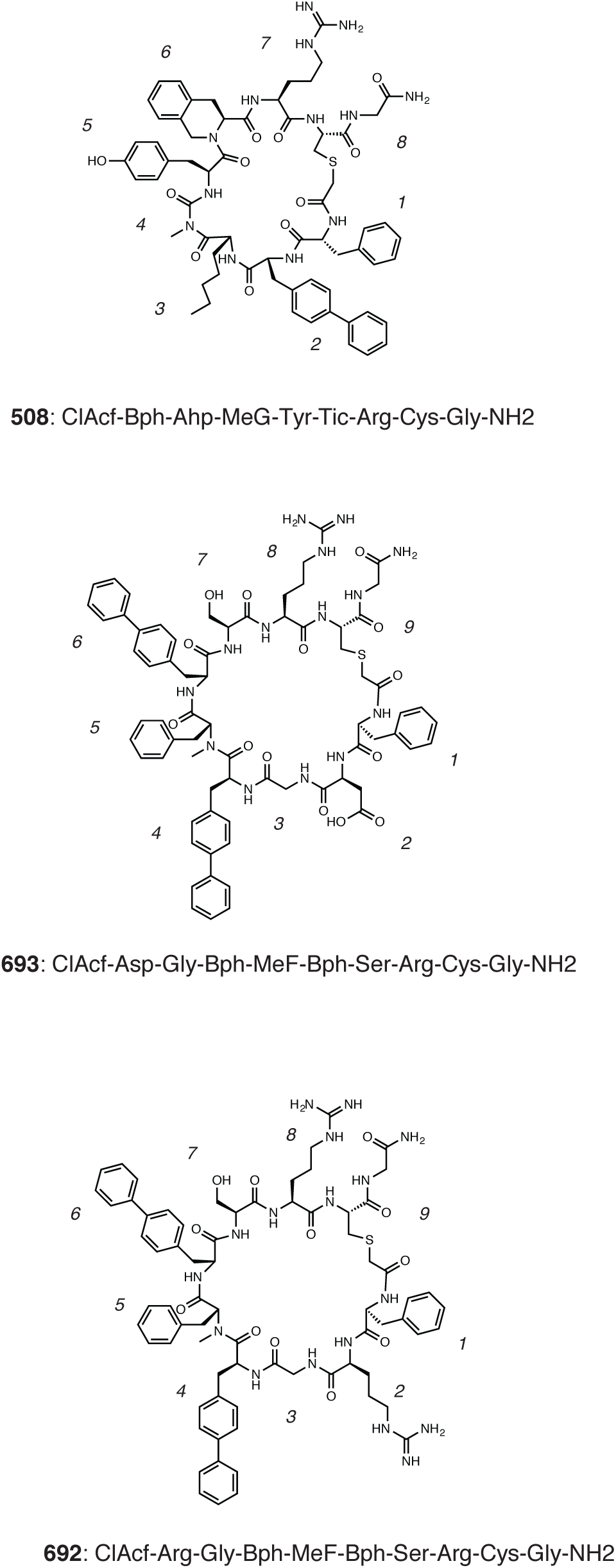
Chemical structures of original hit macrocycles **508**, **692** and **693** identified in the library screen. The sequences of the macrocycles are represented in a linear format using the three letter amino acid codes. Non-natural amino acids are as follows: N-*α*-Methyl-L-phenylalanine (MeF), N-*α*-Methyl-L-glycine (MeG, Sarcosine), (S)-2-Aminoheptanoic acid (Ahp), 4-Phenyl-L-phenylalanine (Bph) and (S)-1,2,3,4-Tetrahydroisoquinoline-3-carboxylic acid (Tic). CIAcf was fixed at the first position and used for cyclization.

**Figure 4-figure supplement 1:**
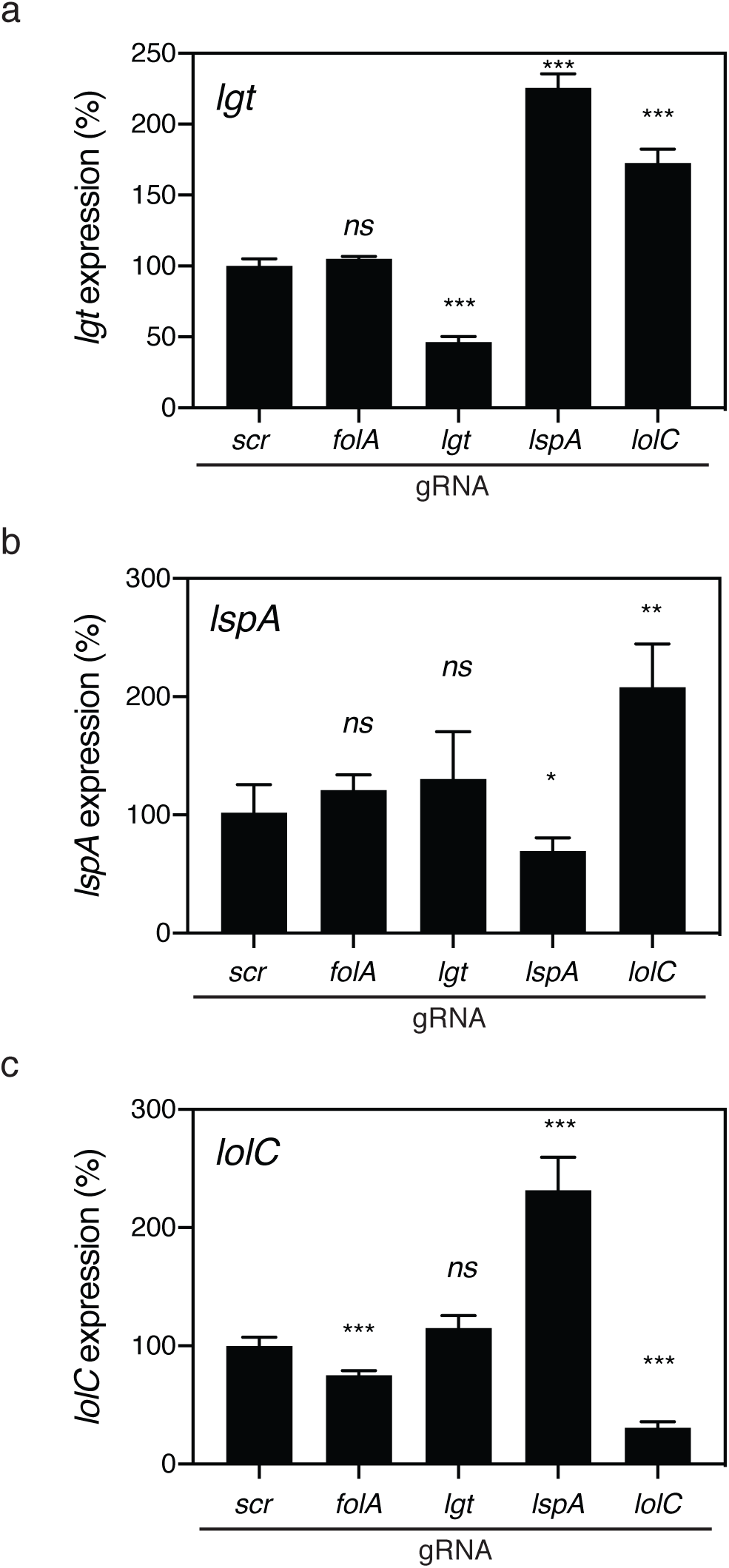
Efficiency of CRISPRi-mediated downregulation of target genes. Total RNA was harvested from *E. coli* BW25113 cells transformed with scrambled (scr) gRNA or gRNA specific for *folA*, *lgt*, *lspA*, and *lolC* and gene expression of *lgt* **(a)**, *lspA* **(b)** and *lolC* **(c)** was measured by RT-qPCR. Relative gene expression of *lgt*, *lspA* and *lolC* were calculated by normalizing to *rpoB* levels using the 2^-^*^ΔΔ^*^CT^ method. Expression levels are graphed after comparison to “scr” gRNA, which was set at 100%. Data are representative of two independent experiments each performed in duplicate (*ns* = not significant, *p < 0.05, **p < 0.01, ***p < 0.001).

**Figure 4-figure supplement 2:**
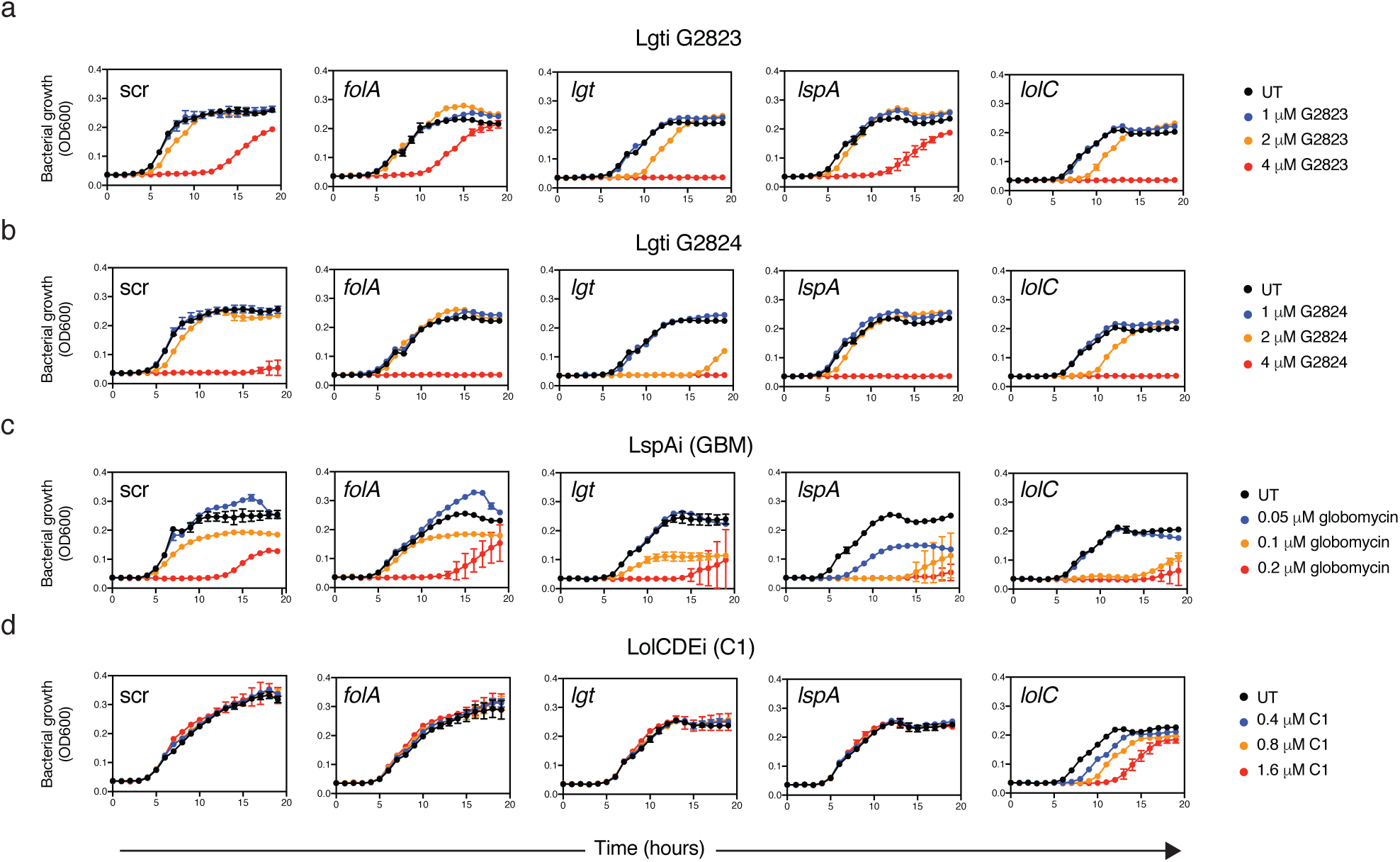
Kinetics of growth inhibition after CRISPRi gRNA induction in *E. coli* BW25113. BW25113 cells expressing either scrambled (scr) gRNA or gRNAs specific to *folA*, *lgt*, *lspA* or *lolC* were treated with **(a)** Lgti G2823, **(b)** Lgti G2824, **(c)** LspAi (GBM) and **(d)** LolCDEi (C1) and bacterial growth was measured by OD_600_. For all inhibitors, three concentrations were tested based on the MIC of each molecule and compared to untreated cells (UT, black) (Lgti = 4, 2 and 1 μM; LspAi = 0.2, 0.1 and 0.05 μM and LolCDEi = 1.6, 0.8 and 0.4 μM). Data are representative of three independent experiments each performed in triplicate.

**Figure 5-figure supplement 1:**
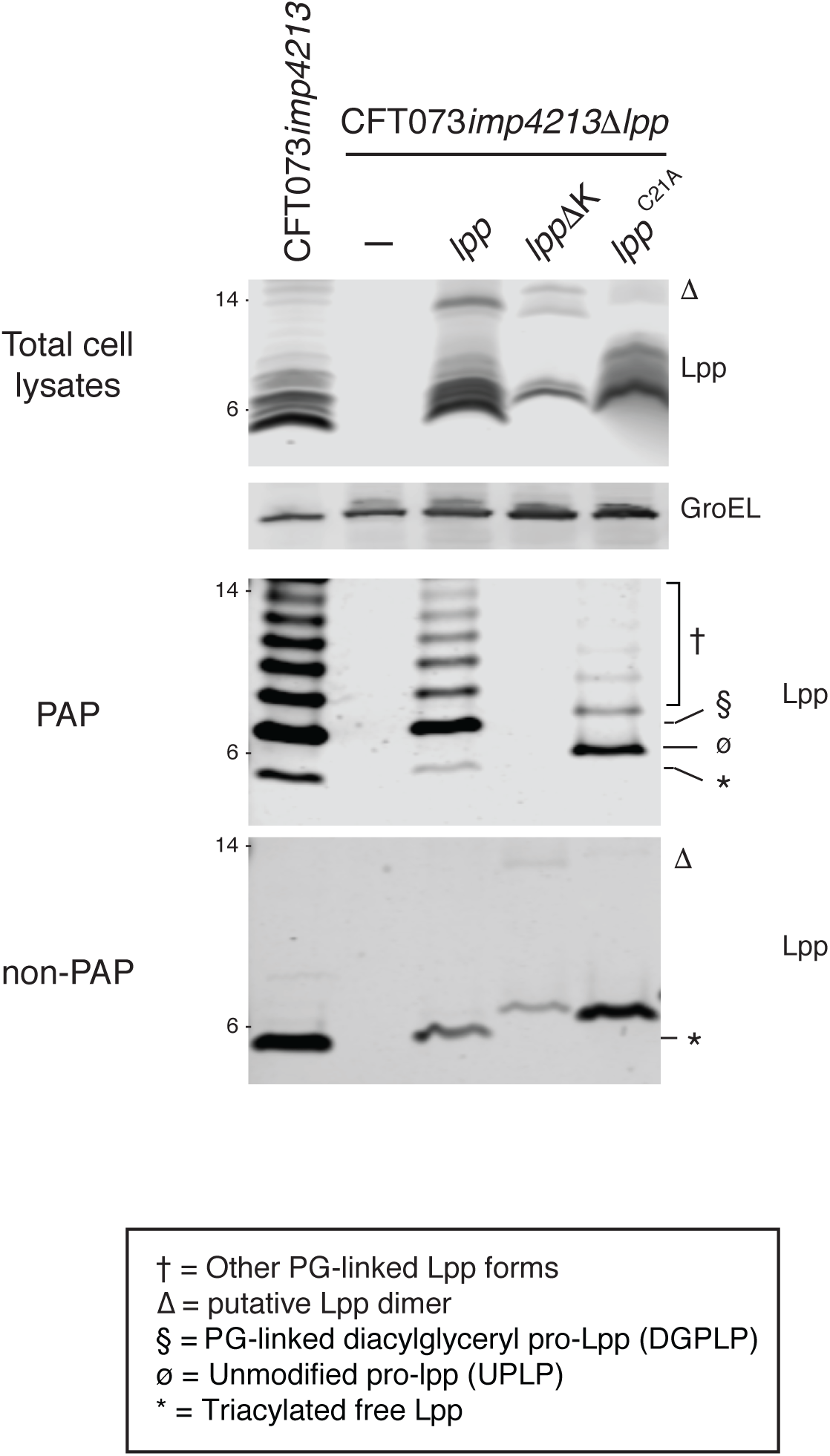
Determination of PG-linkage of WT Lpp, Lpp*Δ*K and Lpp^C21A^. CFT073*imp*4213, CFT073*imp*4213*Δlpp* or CFT073*imp*4213*Δlpp* complemented with WT *lpp*, *lppΔ*K or *lpp*^C21A^ were treated with SDS to isolate PAP and non-PAP fractions. Lpp levels were detected by Western blot analyses in total cell lysates, PAP and non-PAP fractions. The Lpp^C21A^ mutant contains an alanine in place of the conserved cysteine in the lipobox. The *lppΔ*K construct is His-tagged and hence migrates slower on SDS-PAGE. Lpp forms are denoted in the figure (* = triacylated free Lpp; § = PG-linked DGPLP; ø = UPLP; † = other PG-linked Lpp forms; *Δ* = putative Lpp dimer).

**Figure 6-figure supplement 1:**
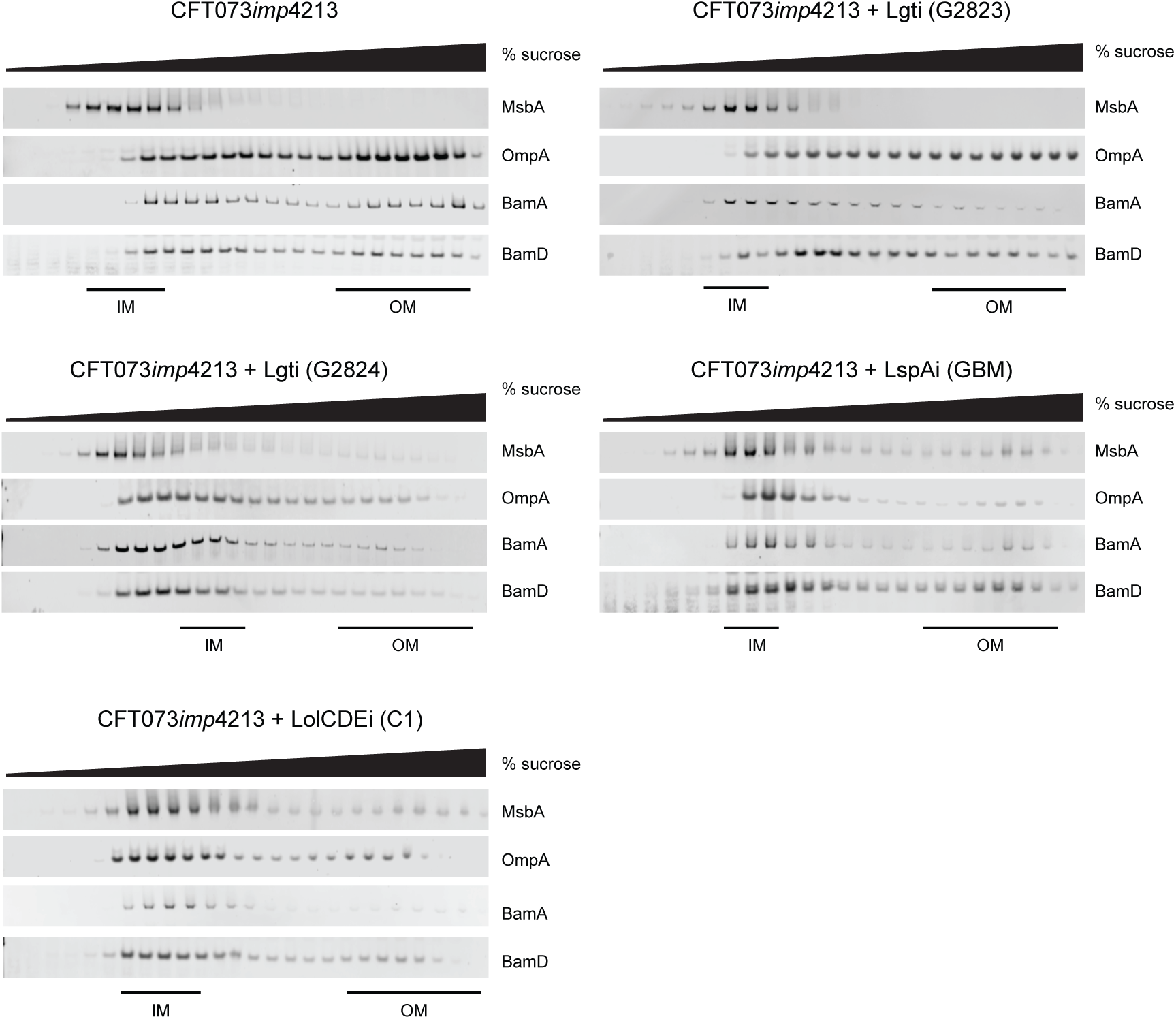
Membrane localization of BamA and BamD in CFT073*imp*4213 cells treated with Lgti (G2823 and G2824), LspAi (GBM) or LolCDEi (C1) for 60 minutes at 1×MIC and subjected to sucrose gradient ultracentrifugation as described in the Methods. Levels of BamA and BamD were detected by Western blot analyses. IM and OM fractions were assigned based on the expression of MsbA and OmpA, respectively, as presented in Figure 6.

**Table S1:** Lgti minimal inhibitory concentrations (MIC) against LolCDE-resistant *E. coli* isolates

| **MG1655*imp4213*** | **MIC (μM)** |  |  |  |  | **Vancomycin** |
| --- | --- | --- | --- | --- | --- | --- |
|  | **Lgti**<br>**(G9066)** | **Lgti**<br>**(G2823)** | **Lgti**<br>**(G2824)** | **LspAi**<br>**(GBM)** | **LolCDEi**<br>**(C2)** |  |
| WT | 3.1 | 4.7 | 3.1 | 0.2 | 2.4 | 0.8 |
| LolC (Q258K) | 6.3 | 3.1 | 3.1 | 0.3 | 75 | 0.8 |
| LolC (N265K) | 3.1 | 6.3 | 3.1 | 0.2 | 37.5 | 1.2 |
| LolD (S43R) | 6.3 | 4.7 | 3.1 | 0.2 | 37.5 | 0.8 |
| LolE (F367L) | 6.3 | 3.1 | 3.1 | 0.3 | 25 | 1.2 |
| LolE (P372L) | 3.1 | 6.3 | 3.1 | 0.2 | 50 | 1.2 |

**Table S2:**
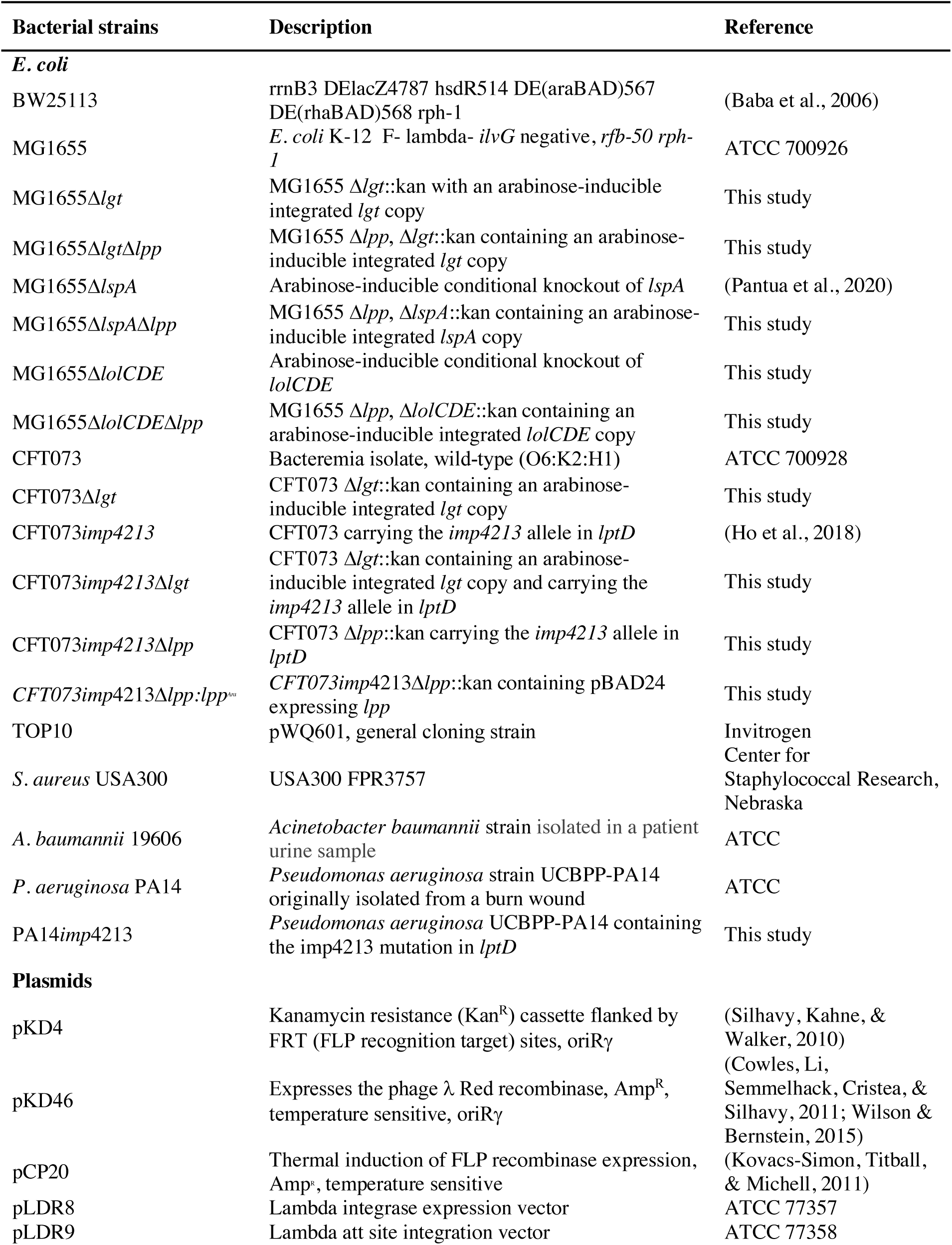
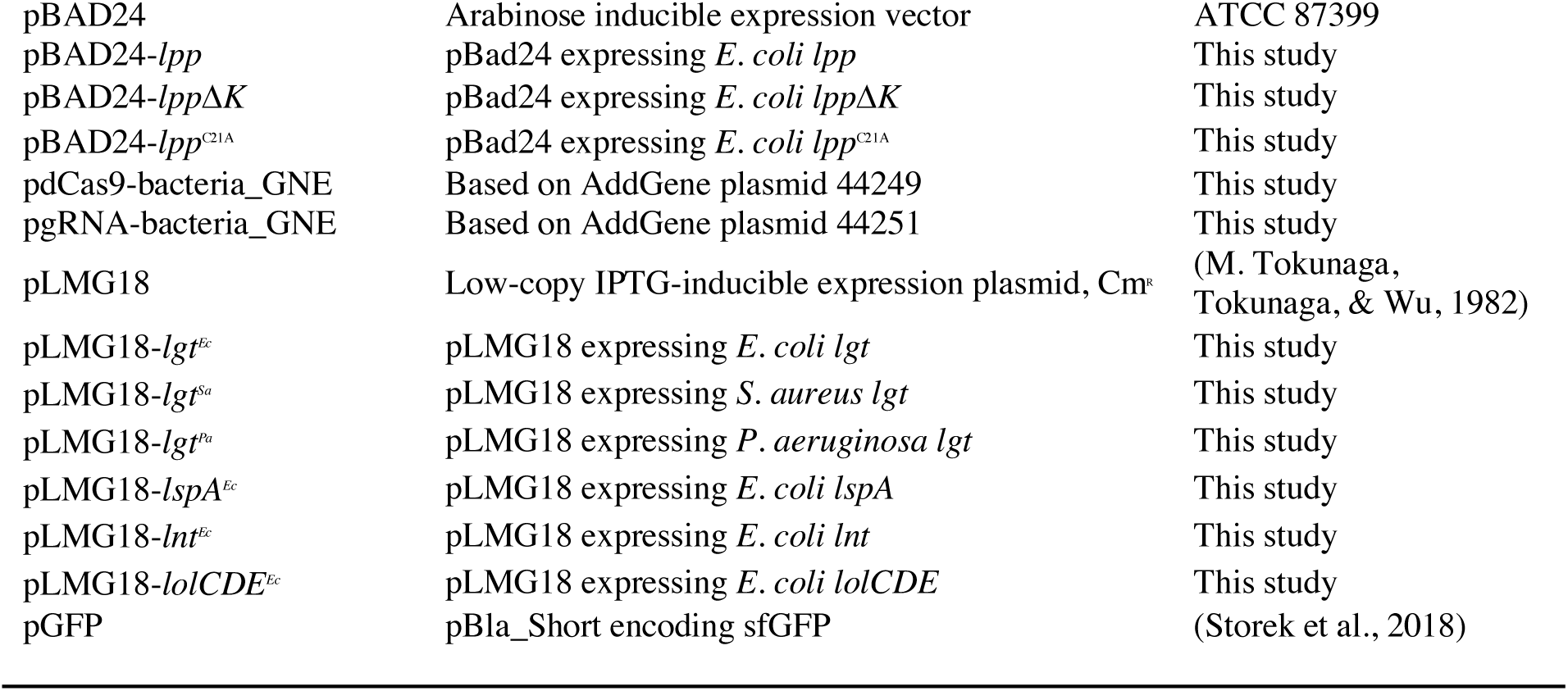
Bacterial strains and plasmids used in this study

---

**Table S3:**
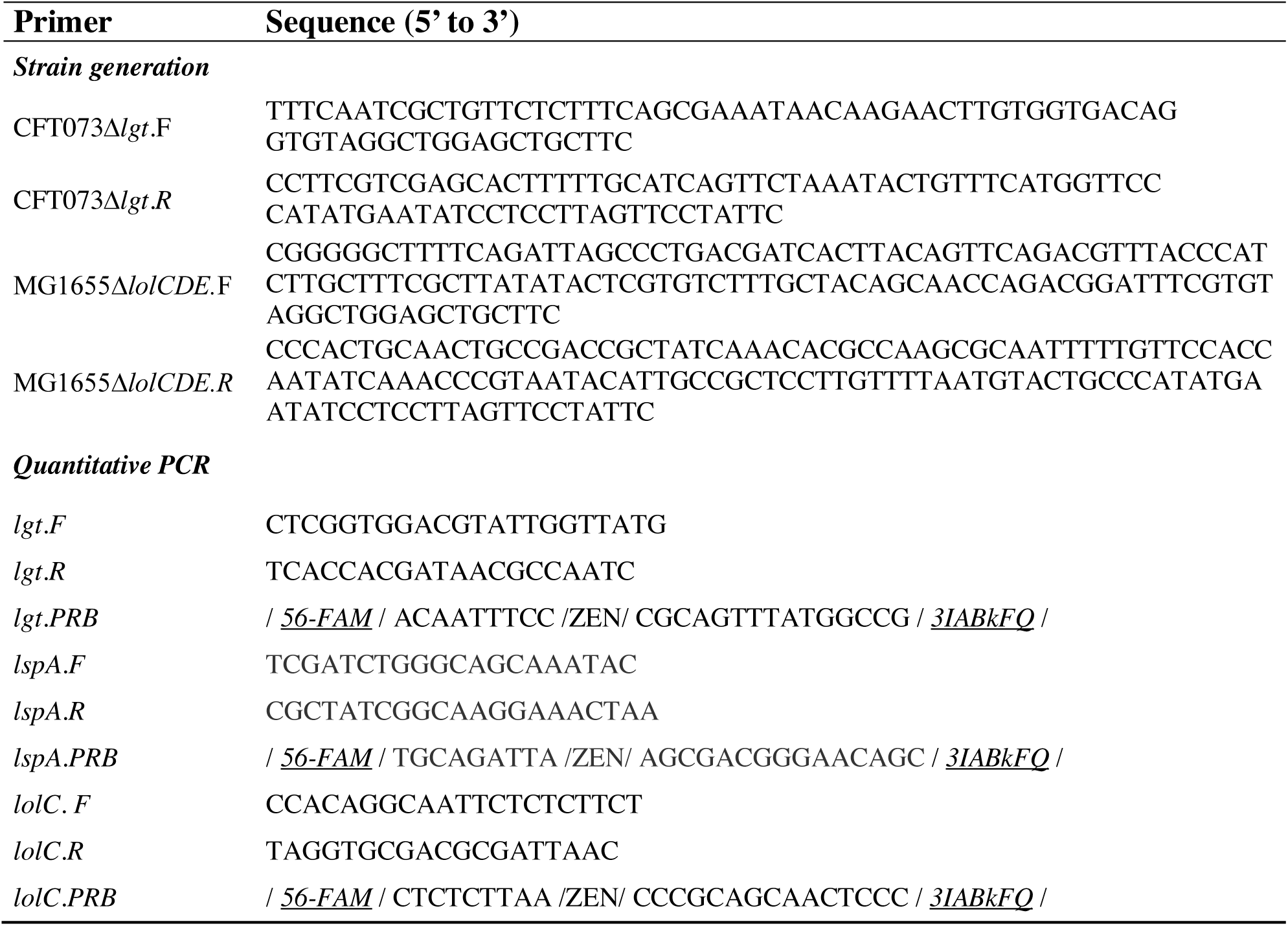
Primers used in this study for strain generation and quantitative PCR

## References

Balibar, C. J., & Grabowicz, M. (2016). Mutant Alleles of lptD Increase the Permeability of Pseudomonas aeruginosa and Define Determinants of Intrinsic Resistance to Antibiotics. Antimicrobial Agents and Chemotherapy, 60(2), 845–854. http://doi.org/10.1128/AAC.01747-15

Bjelić, S., Karshikoff, A., & Jelesarov, I. (2006). Stability and folding/unfolding kinetics of the homotrimeric coiled coil Lpp-56. Biochemistry, 45(29), 8931–8939. http://doi.org/10.1021/bi0608156

Braun, V., & Wolff, H. (1970). The murein-lipoprotein linkage in the cell wall of Escherichia coli. European Journal of Biochemistry / FEBS, 14(2), 387–391.

Caro, F., Place, N. M., & Mekalanos, J. J. (2019). Analysis of lipoprotein transport depletion in Vibrio cholerae using CRISPRi. Proceedings of the National Academy of Sciences of the United States of America, 116(34), 17013–17022. http://doi.org/10.1073/pnas.1906158116

Cascales, E., Bernadac, A., Gavioli, M., Lazzaroni, J.-C., & Lloubes, R. (2002). Pal lipoprotein of Escherichia coli plays a major role in outer membrane integrity. Journal of Bacteriology, 184(3), 754–759.

Chang, T.-W., Lin, Y.-M., Wang, C.-F., & Liao, Y.-D. (2012). Outer membrane lipoprotein Lpp is Gram-negative bacterial cell surface receptor for cationic antimicrobial peptides. The Journal of Biological Chemistry, 287(1), 418–428. http://doi.org/10.1074/jbc.M111.290361

Cherepanov, P. P., & Wackernagel, W. (1995). Gene disruption in Escherichia coli: TcR and KmR cassettes with the option of Flp-catalyzed excision of the antibiotic-resistance determinant. Gene, 158(1), 9–14.

Clavel, T., Germon, P., Vianney, A., Portalier, R., & Lazzaroni, J. C. (1998). TolB protein of Escherichia coli K-12 interacts with the outer membrane peptidoglycan-associated proteins Pal, Lpp and OmpA. Molecular Microbiology, 29(1), 359–367.

Cowles, C. E., Li, Y., Semmelhack, M. F., Cristea, I. M., & Silhavy, T. J. (2011). The free and bound forms of Lpp occupy distinct subcellular locations in Escherichia coli. Molecular Microbiology, 79(5), 1168–1181. http://doi.org/10.1111/j.1365-2958.2011.07539.x

Cross, A. S., Siegel, G., Byrne, W. R., Trautmann, M., & Finbloom, D. S. (1989). Intravenous immune globulin impairs anti-bacterial defences of a cyclophosphamide-treated host. Clinical and Experimental Immunology, 76(2), 159–164.

Datsenko, K. A., & Wanner, B. L. (2000). One-step inactivation of chromosomal genes in Escherichia coli K-12 using PCR products. Proceedings of the National Academy of Sciences of the United States of America, 97(12), 6640–6645. http://doi.org/10.1073/pnas.120163297

Dev, I. K., Harvey, R. J., & Ray, P. H. (1985). Inhibition of prolipoprotein signal peptidase by globomycin. The Journal of Biological Chemistry, 260(10), 5891–5894.

Diao, J., Bouwman, C., Yan, D., Kang, J., Katakam, A. K., Liu, P., et al. (2017). Peptidoglycan Association of Murein Lipoprotein Is Required for KpsD-Dependent Group 2 Capsular Polysaccharide Expression and Serum Resistance in a Uropathogenic Escherichia coli Isolate. mBio, 8(3), e00603–17. http://doi.org/10.1128/mBio.00603-17

Ferrer-Navarro, M., Ballesté-Delpierre, C., Vila, J., & Fàbrega, A. (2016). Characterization of the outer membrane subproteome of the virulent strain Salmonella Typhimurium SL1344. Journal of Proteomics, 146, 141–147. http://doi.org/10.1016/j.jprot.2016.06.032

Filip, C., Fletcher, G., Wulff, J. L., & Earhart, C. F. (1973). Solubilization of the cytoplasmic membrane of Escherichia coli by the ionic detergent sodium-lauryl sarcosinate. Journal of Bacteriology, 115(3), 717–722.

Gan, K., Gupta, S. D., Sankaran, K., Schmid, M. B., & Wu, H. C. (1993). Isolation and characterization of a temperature-sensitive mutant of Salmonella typhimurium defective in prolipoprotein modification. The Journal of Biological Chemistry, 268(22), 16544–16550.

Gan, K., Sankaran, K., Williams, M. G., Aldea, M., Rudd, K. E., Kushner, S. R., & Wu, H. C. (1995). The umpA gene of Escherichia coli encodes phosphatidylglycerol:prolipoprotein diacylglyceryl transferase (lgt) and regulates thymidylate synthase levels through translational coupling. Journal of Bacteriology, 177(7), 1879–1882.

Gerth, K., Irschik, H., Reichenbach, H., & Trowitzsch, W. (1982). The myxovirescins, a family of antibiotics from Myxococcus virescens (Myxobacterales). The Journal of Antibiotics, 35(11), 1454–1459.

Goto, Y., Katoh, T., & Suga, H. (2011). Flexizymes for genetic code reprogramming. Nature Protocols, 6(6), 779–790. http://doi.org/10.1038/nprot.2011.331

Gupta, S. D., Gan, K., Schmid, M. B., & Wu, H. C. (1993). Characterization of a temperature-sensitive mutant of Salmonella typhimurium defective in apolipoprotein N-acyltransferase. The Journal of Biological Chemistry, 268(22), 16551–16556.

Hirota, Y., Suzuki, H., Nishimura, Y., & Yasuda, S. (1977). On the process of cellular division in Escherichia coli: a mutant of E. coli lacking a murein-lipoprotein. Proceedings of the National Academy of Sciences of the United States of America, 74(4), 1417–1420.

Hmelo, L. R., Borlee, B. R., Almblad, H., Love, M. E., Randall, T. E., Tseng, B. S., et al. (2015). Precision-engineering the Pseudomonas aeruginosa genome with two-step allelic exchange. Nature Protocols, 10(11), 1820–1841. http://doi.org/10.1038/nprot.2015.115

Hobb, R. I., Fields, J. A., Burns, C. M., & Thompson, S. A. (2009). Evaluation of procedures for outer membrane isolation from Campylobacter jejuni. Microbiology (Reading, England), 155(Pt 3), 979–988. http://doi.org/10.1099/mic.0.024539-0

Igarashi, M. (2019). New natural products to meet the antibiotic crisis: a personal journey. The Journal of Antibiotics, 72(12), 890–898. http://doi.org/10.1038/s41429-019-0224-6

Inukai, M., Enokita, R., Torikata, A., Nakahara, M., Iwado, S., & Arai, M. (1978a). Globomycin, a new peptide antibiotic with spheroplast-forming activity. I. Taxonomy of producing organisms and fermentation. The Journal of Antibiotics, 31(5), 410–420. http://doi.org/10.7164/antibiotics.31.410

Inukai, M., Nakajima, M., Osawa, M., Haneishi, T., & Arai, M. (1978b). Globomycin, a new peptide antibiotic with spheroplast-forming activity. II. Isolation and physico-chemical and biological characterization. The Journal of Antibiotics, 31(5), 421–425. http://doi.org/10.7164/antibiotics.31.421

Ishizawa, T., Kawakami, T., Reid, P. C., & Murakami, H. (2013). TRAP display: a high-speed selection method for the generation of functional polypeptides. Journal of the American Chemical Society, 135(14), 5433–5440. http://doi.org/10.1021/ja312579u

Jabbour, R. E., Wade, M. M., Deshpande, S. V., Stanford, M. F., Wick, C. H., Zulich, A. W., & Snyder, A. P. (2010). Identification of Yersinia pestis and Escherichia coli strains by whole cell and outer membrane protein extracts with mass spectrometry-based proteomics. Journal of Proteome Research, 9(7), 3647–3655. http://doi.org/10.1021/pr100402y

Kashiwagi, K., Reid, C. P., & Inc, P. (2013). Rapid display method in translational synthesis of peptide.

Kitamura, S., Owensby, A., Wall, D., & Wolan, D. W. (2018). Lipoprotein Signal Peptidase Inhibitors with Antibiotic Properties Identified through Design of a Robust In Vitro HT Platform. Cell Chemical Biology, 25(3), 301–308.e12. http://doi.org/10.1016/j.chembiol.2017.12.011

Kovacs-Simon, A., Titball, R. W., & Michell, S. L. (2011). Lipoproteins of bacterial pathogens. Infection and Immunity, 79(2), 548–561. http://doi.org/10.1128/IAI.00682-10

Kowata, H., Tochigi, S., Kusano, T., & Kojima, S. (2016). Quantitative measurement of the outer membrane permeability in Escherichia coli lpp and tol-pal mutants defines the significance of Tol-Pal function for maintaining drug resistance. The Journal of Antibiotics. http://doi.org/10.1038/ja.2016.50

Leduc, M., Ishidate, K., Shakibai, N., & Rothfield, L. (1992). Interactions of Escherichia coli membrane lipoproteins with the murein sacculus. Journal of Bacteriology, 174(24), 7982–7988.

Lehman, K. M., & Grabowicz, M. (2019). Countering Gram-Negative Antibiotic Resistance: Recent Progress in Disrupting the Outer Membrane with Novel Therapeutics. Antibiotics (Basel, Switzerland), 8(4), 163. http://doi.org/10.3390/antibiotics8040163

Mao, G., Zhao, Y., Kang, X., Li, Z., Zhang, Y., Wang, X., et al. (2016). Crystal structure of E. coli lipoprotein diacylglyceryl transferase. Nature Communications, 7, 10198. http://doi.org/10.1038/ncomms10198

Mathelié-Guinlet, M., Asmar, A. T., Collet, J.-F., & Dufrêne, Y. F. (2020). Lipoprotein Lpp regulates the mechanical properties of the E. coli cell envelope. Nature Communications, 11(1), 1789–11. http://doi.org/10.1038/s41467-020-15489-1

McLeod, S. M., Fleming, P. R., MacCormack, K., McLaughlin, R. E., Whiteaker, J. D., Narita, S.-I., et al. (2015). Small molecule inhibitors of Gram-negative lipoprotein trafficking discovered by phenotypic screening. Journal of Bacteriology, 197(6), JB.02352–14–1082. http://doi.org/10.1128/JB.02352-14

Mizuno, T. (1979). A novel peptidoglycan-associated lipoprotein found in the cell envelope of Pseudomonas aeruginosa and Escherichia coli. Journal of Biochemistry, 86(4), 991–1000.

Mobley, H. L., Green, D. M., Trifillis, A. L., Johnson, D. E., Chippendale, G. R., Lockatell, C. V., et al. (1990). Pyelonephritogenic Escherichia coli and killing of cultured human renal proximal tubular epithelial cells: role of hemolysin in some strains. Infection and Immunity, 58(5), 1281– 1289.

Nakae, T., Ishii, J., & Tokunaga, M. (1979). Subunit structure of functional porin oligomers that form permeability channels in the other membrane of Escherichia coli. The Journal of Biological Chemistry, 254(5), 1457–1461.

Narita, S.-I. (2011). ABC transporters involved in the biogenesis of the outer membrane in gram-negative bacteria. Bioscience, Biotechnology, and Biochemistry, 75(6), 1044–1054. http://doi.org/10.1271/bbb.110115

Narita, S.-I., & Tokuda, H. (2010). Sorting of bacterial lipoproteins to the outer membrane by the Lol system. Methods in Molecular Biology (Clifton, NJ), 619(Chapter 7), 117–129. http://doi.org/10.1007/978-1-60327-412-8_7

Narita, S.-I., & Tokuda, H. (2011). Overexpression of LolCDE allows deletion of the Escherichia coli gene encoding apolipoprotein N-acyltransferase. Journal of Bacteriology, 193(18), 4832– 4840. http://doi.org/10.1128/JB.05013-11

Neidhardt, F. C. (1996). Chemical composition of Escherichia coli. In Escherichia coli and Salmonella: cellular and molecular biology (2nd ed., pp. 1035–1063). Washington D.C.

Nickerson, N. N., Jao, C. C., Xu, Y., Quinn, J., Skippington, E., Alexander, M. K., et al. (2018). A Novel Inhibitor of the LolCDE ABC Transporter Essential for Lipoprotein Trafficking in Gram-Negative Bacteria. Antimicrobial Agents and Chemotherapy, 62(4), e02151–17. http://doi.org/10.1128/AAC.02151-17

Noland, C. L., Kattke, M. D., Diao, J., Gloor, S. L., Pantua, H., Reichelt, M., et al. (2017). Structural insights into lipoprotein N-acylation by Escherichia coli apolipoprotein N-acyltransferase. Proceedings of the National Academy of Sciences of the United States of America, 114(30), E6044–E6053. http://doi.org/10.1073/pnas.1707813114

Olatunji, S., Yu, X., Bailey, J., Huang, C.-Y., Zapotoczna, M., Bowen, K., et al. (2020). Structures of lipoprotein signal peptidase II from Staphylococcus aureus complexed with antibiotics globomycin and myxovirescin. Nature Communications, 11(1), 140–11. http://doi.org/10.1038/s41467-019-13724-y

Pailler, J., Aucher, W., Pires, M., & Buddelmeijer, N. (2012). Phosphatidylglycerol::prolipoprotein diacylglyceryl transferase (Lgt) of Escherichia coli has seven transmembrane segments, and its essential residues are embedded in the membrane. Journal of Bacteriology, 194(9), 2142–2151. http://doi.org/10.1128/JB.06641-11

Palmer, A. C., & Kishony, R. (2014). Opposing effects of target overexpression reveal drug mechanisms. Nature Communications, 5, 4296. http://doi.org/10.1038/ncomms5296

Pantua, H., Skippington, E., braun, M.-G., Noland, C. L., Diao, J., peng, Y., et al. (2020). Unstable Mechanisms of Resistance to Inhibitors of Escherichia coli Lipoprotein Signal Peptidase. mBio, 11(5), VMBF–0016–2015. http://doi.org/10.1128/mBio.02018-20

Peters, J. M., Colavin, A., Shi, H., Czarny, T. L., Larson, M. H., Wong, S., et al. (2016). A Comprehensive, CRISPR-based Functional Analysis of Essential Genes in Bacteria. Cell, 165(6), 1493–1506. http://doi.org/10.1016/j.cell.2016.05.003

Qi, L. S., Larson, M. H., Gilbert, L. A., Doudna, J. A., Weissman, J. S., Arkin, A. P., & Lim, W. A. (2013). Repurposing CRISPR as an RNA-guided platform for sequence-specific control of gene expression. Cell, 152(5), 1173–1183. http://doi.org/10.1016/j.cell.2013.02.022

Robichon, C., Vidal-Ingigliardi, D., & Pugsley, A. P. (2005). Depletion of apolipoprotein N-acyltransferase causes mislocalization of outer membrane lipoproteins in Escherichia coli. The Journal of Biological Chemistry, 280(2), 974–983. http://doi.org/10.1074/jbc.M411059200

Rojas, E. R., Billings, G., Odermatt, P. D., Auer, G. K., Zhu, L., Miguel, A., et al. (2018). The outer membrane is an essential load-bearing element in Gram-negative bacteria. Nature, 559(7715), 617–621. http://doi.org/10.1038/s41586-018-0344-3

Rossiter, S. E., Fletcher, M. H., & Wuest, W. M. (2017). Natural Products as Platforms To Overcome Antibiotic Resistance. Chemical Reviews, 117(19), 12415–12474. http://doi.org/10.1021/acs.chemrev.7b00283

Rousset, F., Cui, L., Siouve, E., Becavin, C., Depardieu, F., & Bikard, D. (2018). Genome-wide CRISPR-dCas9 screens in E. coli identify essential genes and phage host factors. PLoS Genetics, 14(11), e1007749. http://doi.org/10.1371/journal.pgen.1007749

Ruiz, N., Falcone, B., Kahne, D., & Silhavy, T. J. (2005). Chemical conditionality: a genetic strategy to probe organelle assembly. Cell, 121(2), 307–317. http://doi.org/10.1016/j.cell.2005.02.014

Sankaran, K., & Wu, H. C. (1994). Lipid modification of bacterial prolipoprotein. Transfer of diacylglyceryl moiety from phosphatidylglycerol. The Journal of Biological Chemistry, 269(31), 19701–19706.

Schindelin, J., Arganda-Carreras, I., Frise, E., Kaynig, V., Longair, M., Pietzsch, T., et al. (2012). Fiji: an open-source platform for biological-image analysis. Nature Methods, 9(7), 676–682. http://doi.org/10.1038/nmeth.2019

Schlesinger, M. J. (1992). Lipid Modifications of Proteins. CRC Press.

Shu, W., Liu, J., Ji, H., & Lu, M. (2000). Core structure of the outer membrane lipoprotein from Escherichia coli at 1.9 A resolution. Journal of Molecular Biology, 299(4), 1101–1112. http://doi.org/10.1006/jmbi.2000.3776

Silhavy, T. J., Kahne, D., & Walker, S. (2010). The bacterial cell envelope. Cold Spring Harbor Perspectives in Biology, 2(5), a000414–a000414. http://doi.org/10.1101/cshperspect.a000414

Singh, W., Bilal, M., McClory, J., Dourado, D., Quinn, D., Moody, T. S., et al. (2019). Mechanism of Phosphatidylglycerol Activation Catalyzed by Prolipoprotein Diacylglyceryl Transferase. The Journal of Physical Chemistry. B, 123(33), 7092–7102. http://doi.org/10.1021/acs.jpcb.9b04227

Stoll, H., Dengjel, J., Nerz, C., & Götz, F. (2005). Staphylococcus aureus deficient in lipidation of prelipoproteins is attenuated in growth and immune activation. Infection and Immunity, 73(4), 2411–2423. http://doi.org/10.1128/IAI.73.4.2411-2423.2005

Storek, K. M., Auerbach, M. R., Shi, H., Garcia, N. K., Sun, D., Nickerson, N. N., et al. (2018). Monoclonal antibody targeting the β-barrel assembly machine of Escherichia coli is bactericidal. Proceedings of the National Academy of Sciences of the United States of America, 115(14), 3692–3697. http://doi.org/10.1073/pnas.1800043115

Storek, K. M., Chan, J., Vij, R., Chiang, N., Lin, Z., Bevers, J., et al. (2019). Massive antibody discovery used to probe structure-function relationships of the essential outer membrane protein LptD. eLife, 8, 3002. http://doi.org/10.7554/eLife.46258

Suzuki, H., Nishimura, Y., Yasuda, S., Nishimura, A., Yamada, M., & Hirota, Y. (1978). Murein-lipoprotein of Escherichia coli: a protein involved in the stabilization of bacterial cell envelope. Molecular & General Genetics : MGG, 167(1), 1–9.

Suzuki, M., Hara, H., & Matsumoto, K. (2002). Envelope disorder of Escherichia coli cells lacking phosphatidylglycerol. Journal of Bacteriology, 184(19), 5418–5425. http://doi.org/10.1128/jb.184.19.5418-5425.2002

Tokunaga, M., Tokunaga, H., & Wu, H. C. (1982). Post-translational modification and processing of Escherichia coli prolipoprotein in vitro. Proceedings of the National Academy of Sciences of the United States of America, 79(7), 2255–2259. http://doi.org/10.1073/pnas.79.7.2255

Vogeley, L., Arnaout, El, T., Bailey, J., Stansfeld, P. J., Boland, C., & Caffrey, M. (2016). Structural basis of lipoprotein signal peptidase II action and inhibition by the antibiotic globomycin. Science (New York, N.Y.), 351(6275), 876–880. http://doi.org/10.1126/science.aad3747

Whitfield, C., Hancock, R. E., & Costerton, J. W. (1983). Outer membrane protein K of Escherichia coli: purification and pore-forming properties in lipid bilayer membranes. Journal of Bacteriology, 156(2), 873–879.

Wilson, M. M., & Bernstein, H. D. (2015). Surface-Exposed Lipoproteins: An Emerging Secretion Phenomenon in Gram-Negative Bacteria. Trends in Microbiology. http://doi.org/10.1016/j.tim.2015.11.006

Xiao, Y., Gerth, K., Müller, R., & Wall, D. (2012). Myxobacterium-produced antibiotic TA (myxovirescin) inhibits type II signal peptidase. Antimicrobial Agents and Chemotherapy, 56(4), 2014–2021. http://doi.org/10.1128/AAC.06148-11

Yakushi, T., Tajima, T., Matsuyama, S., & Tokuda, H. (1997a). Lethality of the covalent linkage between mislocalized major outer membrane lipoprotein and the peptidoglycan of Escherichia coli. Journal of Bacteriology, 179(9), 2857–2862.

Yakushi, T., Tajima, T., Matsuyama, S., & Tokuda, H. (1997b). Lethality of the covalent linkage between mislocalized major outer membrane lipoprotein and the peptidoglycan of Escherichia coli. Journal of Bacteriology, 179(9), 2857–2862.

Yem, D. W., & Wu, H. C. (1978). Physiological characterization of an Escherichia coli mutant altered in the structure of murein lipoprotein. Journal of Bacteriology, 133(3), 1419–1426.

Zhang, W. Y., & Wu, H. C. (1992). Alterations of the carboxyl-terminal amino acid residues of Escherichia coli lipoprotein affect the formation of murein-bound lipoprotein. The Journal of Biological Chemistry, 267(27), 19560–19564.

Zhang, W. Y., Inouye, M., & Wu, H. C. (1992). Neither lipid modification nor processing of prolipoprotein is essential for the formation of murein-bound lipoprotein in Escherichia coli. The Journal of Biological Chemistry, 267(27), 19631–19635.

Zwiebel, L. J., Inukai, M., Nakamura, K., & Inouye, M. (1981). Preferential selection of deletion mutations of the outer membrane lipoprotein gene of Escherichia coli by globomycin. Journal of Bacteriology, 145(1), 654–656.

## References

Baba, T., Ara, T., Hasegawa, M., Takai, Y., Okumura, Y., Baba, M., et al. (2006). Construction of Escherichia coli K-12 in-frame, single-gene knockout mutants: the Keio collection. Molecular Systems Biology, 2(1), 2006.0008. http://doi.org/10.1038/msb4100050

Cowles, C. E., Li, Y., Semmelhack, M. F., Cristea, I. M., & Silhavy, T. J. (2011). The free and bound forms of Lpp occupy distinct subcellular locations in Escherichia coli. Molecular Microbiology, 79(5), 1168– 1181. http://doi.org/10.1111/j.1365-2958.2011.07539.x

Ho, H., Miu, A., Alexander, M. K., Garcia, N. K., Oh, A., Zilberleyb, I., et al. (2018). Structural basis for dual-mode inhibition of the ABC transporter MsbA. Nature, 557(7704), 196–201. http://doi.org/10.1038/s41586-018-0083-5

Pantua, H., Skippington, E., Braun, M.-G., Noland, C. L., Diao, J., Peng, Y., et al. (2020). Unstable Mechanisms of Resistance to Inhibitors of Escherichia coli Lipoprotein Signal Peptidase. mBio, 11(5), VMBF–0016–2015. http://doi.org/10.1128/mBio.02018-20

